# A Mechanism Underpinning the Bioenergetic Metabolism-Regulating Function of Gold Nanocatalysts

**DOI:** 10.1101/2023.05.08.539856

**Authors:** Zixin Wang, Alexandre Henriques, Laura Rouvière, Noëlle Callizot, Lin Tan, Michael T. Hotchkin, Rodrigue Rossignol, Mark G. Mortenson, Karen S. Ho, Hui Wang

## Abstract

Bioenergetic deficits, such as mitochondrial impairments and dysfunction in glucose metabolism, have been identified as significant contributors to neurodegenerative diseases. Nevertheless, identifying safe and effective means to address intracellular bioenergetic deficits remains a significant challenge. This work provides mechanistic insights into the bioenergetic metabolism-regulating function of a suspension of gold (Au) nanocrystals, referred to as CNM-Au8®, that are synthesized electrochemically in the absence of any surface-capping organic ligands. When neurons are subjected to excitotoxic stressors or toxic peptides, treatment of neurons with CNM-Au8 results in dose-dependent neuronal survival and preservation of neurite networks across multiple neuronal subtypes. CNM-Au8 efficiently catalyzes the conversion of an energetic co-factor, nicotinamide adenine dinucleotide hydride (NADH), into its oxidized, dehydrogenated counterpart (NAD^+^), which triggers an increase in energy production in the form of adenosine triphosphate (ATP). Detailed kinetic measurements reveal that CNM-Au8-catalyzed NADH oxidation obeys Michaelis-Menten kinetics and exhibits pH-dependent kinetic profiles. CNM-Au8 functions as an NADH-dehydrogenase-mimicking nanozyme that effectively regulates intracellular bioenergetic metabolism. We further utilize photoexcited charge carriers and photothermal transduction, which can be generated through optical excitations of the plasmonic electron oscillations or the interband electronic transitions in CNM-Au8, as unique leverages to modulate reaction kinetics. Benefiting from their bio-compatibility, blood-brain barrier penetrance, tunable optical properties, and enzyme-mimicking functions, CNM-Au8 nanocrystals with deliberately tailored structures and surfactant-free clean surfaces hold great promise for developing next-generation therapeutic agents for neurodegenerative diseases.

---

The high energy demands of the brain make this organ inherently vulnerable to metabolic perturbations that impact adenosine triphosphate (ATP) synthesis and utilization.^1^ As humans age, metabolic inefficiencies increase in number and degree, putting vulnerable neurons at risk of bioenergetic failure.^2^ Several independent lines of evidence indicate that neurodegenerative diseases, such as Alzheimer’s Disease (AD), Parkinson’s disease (PD), and many others, may result as a manifestation of brain bioenergetic failure associated with aging.^1, 3^ Improving cellular bioenergetic metabolism may therefore be a promising target for developing novel neurodegenerative disease-modifying therapies.

Nicotinamide adenine dinucleotide (NAD) is a coenzyme that plays crucial roles in bioenergetic metabolism in all living cells. NAD exists in two forms: a reduced form, NADH, and an oxidized form, NAD^+^. A deficient level of NAD^+^ has been considered a hallmark of bioenergetic deficit as well as a potential therapeutic target for many mitochondrial and neurodegenerative diseases. For example, boosting NAD^+^ levels using genetic or dietary means significantly improves neuronal function, cognition, and survival in AD model organisms.^4–7^ Therapeutic benefits of NAD^+^ have also been documented in models of PD,^8, 9^ amyotrophic lateral sclerosis (ALS),^10, 11^ and multiple sclerosis (MS).^12, 13^ A study of prion disease demonstrated that highly toxic, misfolded prion proteins induced severe, neuron-specific NAD^+^ depletion, followed by decreased intracellular ATP levels.^14^ Replenishment of NAD^+^ to media of prion-treated neuroblastoma cells or provided by intracerebellar injection to prion-protein infected mice resulted in significant rescue of neuroblastoma and hippocampal neurons, respectively, from apoptosis.^14^ In contrast, attempts to reduce prion protein levels by activating proteolytic or autophagic pathways had negligible effect on neuronal survival.^14^

Nanoparticles made of noble metal elements, such as Au, platinum (Pt), and ruthenium (Ru), can exhibit remarkable catalytic activities toward various biologically relevant chemical reactions, acting as potent, stable enzyme mimetics.^15–21^ Colloidal Au nanoparticles can catalyze the aerobic oxidation of NADH to produce NAD^+^ under physiologic reaction conditions in an aqueous environment,^22^ potentially endowing them with direct therapeutic relevance for a number of neurodegenerative diseases. When harnessing the intrinsic catalytic activity of Au nanoparticles for therapeutic applications, clean nanoparticle surfaces with abundant, fully exposed, and easily accessible active sites but that are free of any toxic capping ligands are highly desired. However, traditional chemical reduction synthesis of colloidal Au nanoparticles relies critically on the use of surface-capping organic ligands, such as amphiphilic surfactant molecules, to guide the nanocrystal growth and stabilize the colloidal suspensions.^23–25^ These organic ligands residing on nanoparticle surfaces, which remain difficult and costly to remove in numerous cases,^26, 27^ may not only cause cytotoxicity issues^28–31^ but also lead to compromised catalytic activities.^27, 32–34^ CNM-Au8®is a pharmaceutical grade suspension of Au nanocrystals synthesized using a patented electrochemical method in the absence of any surface-capping organic ligands (CNM-Au8® is a federally-regulated trademark of Clene Nanomedicine, Inc).^35, 36^ CNM-Au8 is the first Au nanocrystal suspension being developed as a therapeutic drug with evidence of remyelinating and neuroprotective activities. CNM-Au8 is administered orally, penetrates the blood brain barrier, and has a clean safety, tolerability, and toxicology profile as demonstrated by multiple, recently completed Phase 2 clinical trials, a First-in-Human study, and standard ICH M3 (R2) animal chronic toxicity studies. Phase 2 proof-of-concept studies of CNM-Au8 for the treatment of MS, ALS, and PD are registered in Clinicaltrials.gov (REPAIR-MS: NCT03993171 and VISIONARY-MS: NCT03536559 and NCT04626921; RESCUE-ALS: NCT04098406, HEALEY-ALS Platform trial: NCT04414345, and REPAIR-PD: NCT03815916).

Our previous preclinical studies on the remyelination activities have revealed that CNM-Au8 exhibits intrinsic capabilities to promote NADH oxidation, to induce the differentiation of oligodendrocyte precursor cells into myelinating oligodendrocytes, and to improve remyelination in two independent demyelinating animal models.^37^ These promising preclinical results motivated us to further investigate the neuroprotective properties of CNM-Au8, as detailed below. The neuroprotective function of CNM-Au8 arises from the aerobic oxidation of NADH catalyzed by colloidal Au nanocrystals, which is found to obey the Michaelis-Menten kinetic model. The kinetic results reported in this work indicate that each nanocrystal in CNM-Au8 behaves as a NADH dehydrogenase-mimicking nanozyme that regulates the bioenergetic metabolism in living cells. We further demonstrate that photoexcited charge carriers and photothermal transduction resulting from optical excitations and decay of the electronic interband transitions and the localized plasmon resonances in Au nanocrystals can be deliberately harnessed as unique leverages to modulate the kinetic enhancement of the Au-catalyzed NADH oxidation reactions.

## RESULTS AND DISCUSSION

### Neuroprotective Function of CNM-Au8

Neural sensitivity to glutamate excitotoxicity has been considered a significant contributory factor to neurodegeneration.^38^ We used validated *in vitro* neuroprotection assays to assess the neuroprotective efficacies of CNM-Au8 treatment against a range of different primary neural-glial co-cultures. As shown in Figure 1A and 1B, treatment of rat primary motor neurons with CNM-Au8 resulted in dose-responsive, statistically significant survival and preservation of neurite lengths following glutamate excitotoxicity in comparison to the treatment with the vehicle control (6.5 mM NaHCO3 without CNM-Au8). We chose riluzole as a comparator because it is capable of blocking glutamate neurotransmission^39^ and is one of three Food and Drug Administration (FDA)-approved drugs for the motor neuron degenerative disease, ALS. CNM-Au8 treatment outperformed riluzole in protecting motor neurons from exogenous glutamate. In addition to primary motor neuron cultures, CNM-Au8 also effectively protected primary hippocampal and cortical neurons from glutamate excitotoxicity in a dose-dependent manner (Figure S1 in the Supporting Information). Improvements in both neuron survival counts, preservation of synapses, and neurite network lengths were achieved upon CNM-Au8 treatment. These experimental results clearly demonstrated that CNM-Au8 exerted neuroprotective activity over central nervous system cells in a manner agnostic to neuronal subtype.

**Figure 1.**
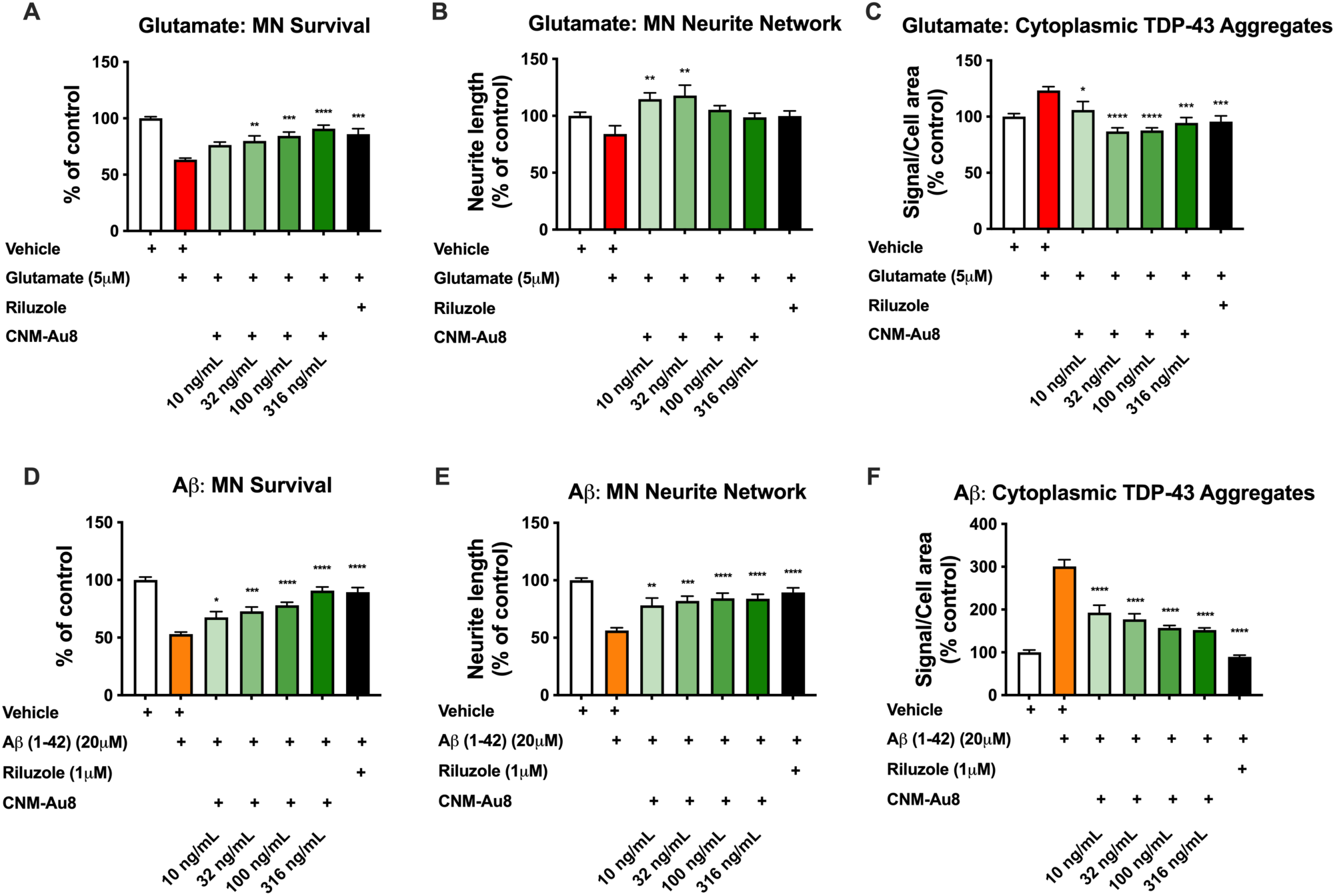
CNM-Au8 treatment increases neuronal survival and protects neurite networks from glutamate challenge and from amyloid-beta oligomer toxicity in vitro. Primary rat E15 spinal motor neurons (MN) were cultured for 11 days then treated with vehicle, CNM-Au8, or riluzole (1µM, 1 h). Glutamate (5 µM) (Panels A-C) or amyloid-beta (1-42) oligomers (20 µM) (Panels D-F) were added to the cultures to induce neuronal death. Cells were fixed and stained with anti-neurofilament to quantitate neuron survival (A, D) and neurite length (B, E), and co-stained with anti-TDP-43 and DAPI to quantitate cytoplasmic (non-DAPI-overlapping) TDP-43 signal (C, F). Six replicates were performed per condition. Group means +/-SEM. * p <0.05; ** p < 0.01; *** p < 0.001; **** p< 0.0001; treatment *vs.* vehicle, one way ANOVA corrected for multiple comparisons.

We have previously shown that CNM-Au8 elevates NAD^+^ levels in primary mesencephalic co-cultures that include dopaminergic neurons.^37^ Treatment of primary rat mesencephalic cultures with 6-hydroxydopamine (6-OHDA) causes accumulation of α-syn aggregates, which can be quantified using immunohistochemistry.^40^ We used this assay as a means of investigating the effect of CNM-Au8 on aggregation of α-syn in tyrosine hydroxylase-expressing (TH+) dopaminergic cells. We observed a dose-dependent reduction in the fraction of TH+ neurons containing α-syn aggregates upon treatment with CNM-Au8 (Figure S2 in the Supporting Information). Similarly, we also studied whether CNM-Au8 affects the mislocalization and aggregation of TDP-43, a protein involved in ALS and also found in a dementia that presents similarly to AD called limbic-predominant age-related TDP-43 encephalopathy.^41, 42^ As shown in Figure 1C, CNM-Au8 treatment led to the significant reduction of mis-localized, cytoplasmic TDP-43 aggregates in motor neurons exposed to glutamate.

The accumulation of amyloid plaques, which consist primarily of extracellular brain deposits of amyloid-beta (Aβ) peptide, is not only a hallmark of AD but also thought to be intimately involved in its pathology. Application of Aβ peptide (1-42 peptide oligomers) to rat primary motor neurons causes motor neuron death, a retraction of neurite processes of the surviving motor neurons, and accumulation of mislocalized TDP-43 in the cytoplasm.^43^ We utilized this assay to investigate the neuroprotective properties of CNM-Au8 when sensitive neurons are exposed to Aβ peptide. Treatment with CNM-Au8 effectively protected the motor neurons against the effects of Aβ peptide on neuron viability (Figure 1D), neurite process lengths (Figure 1E), and aberrant TDP-43 localization (Figure 1F). Based on these results, we conclude that CNM-Au8 exerts pan-protective effects against a toxic protein of direct relevance to neurodegenerative disease pathophysiology.

The neuroprotective function of CNM-Au8 is believed to be intimately tied to Au-catalyzed conversion of NADH to NAD^+^ in the intracellular environment, which could form an important part of the mechanism underpinning the therapeutic effects of CNM-Au8.^37^ Because of the role of NAD^+^ in bioenergetic metabolism regulation, an increase in the intracellular NAD^+^ pool promoted by CNM-Au8 would be expected to result in higher intracellular levels of ATP, which was confirmed by the results shown in Figure 2. In this case, nanocrystals of CNM-Au8 served as an efficient nanocatalyst that kinetically boosted the aerobic oxidation of NADH to produce NAD^+^, effectively regulating the bioenergetic metabolism and protecting the neurons from degeneration. Here we studied the detailed kinetic features of the CNM-Au8-catalyzed NADH oxidation reactions, which enabled us to gain mechanistic insights into the origin of the bioenergetic metabolism-regulating function of Au nanocatalysts.

**Figure 2.**
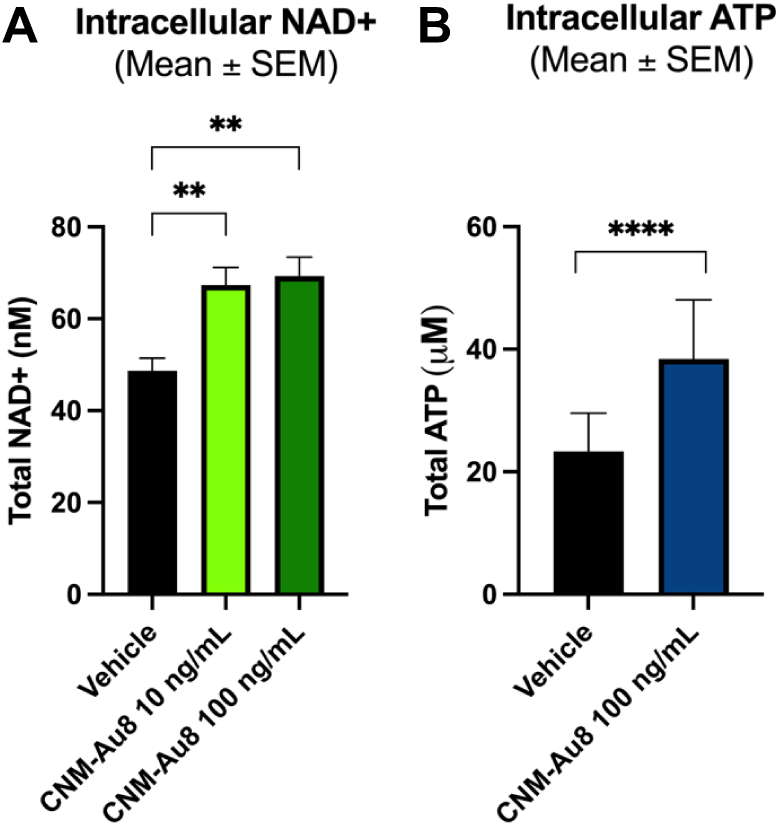
CNM-Au8 treatment increases intracellular levels of NAD+ and ATP. (A) Primary rat E15 mesencephalic co-cultures were grown for four days and then treated with vehicle or CNM-Au8 for 48h. Cells were then lysed and intracellular levels of NAD+ were quantitated. Six replicates were performed per condition. (B) Human glial M03.13 cells were cultured with vehicle or CNM-Au8 for 72h. Total intracellular ATP was quantitated in cell lysates by luciferase assay. Three replicates were performed per condition. Group means +/-SEM. * p <0.05; ** p < 0.01; *** p < 0.001; **** p< 0.0001; treatment vs. vehicle, two-tailed t-test.

### Michaelis-Menten Kinetics of CNM-Au8-Catalyzed NADH Oxidation

As schematically illustrated in Figure 3A, the complete redox reaction catalyzed by CNM-Au8 involved the participation of NADH, H^+^, and the molecular O2 dissolved in water, with NAD^+^ and H2O serving as the products of the oxidation and reduction half reactions, respectively. The CNM-Au8 nanocrystals exhibited a quasi-spherical morphology and a multiply twinned crystalline structure (Figure 3B), with particle diameters narrowly distributed around 11.3 nm (Figure 3C). Although synthesized in a unique reaction environment free of any organic capping ligands, the CNM-Au8 nanocrystals exhibited excellent colloidal stability in aqueous reaction environments over a broad range of pHs because their surfaces were protected by physiosorbed negatively anions. The crystalline and surface structures of CNM-Au8 were characterized in greater detail in a previously published work.^37^ Here we further characterized the surface atomic coordinations and measured the specific surface areas of CNM-Au8 nanocrystals using a cyclic voltammetry (CV)-based electrochemical oxide stripping assay.^44–49^ During the anodic potential sweep, the Au surface atoms were oxidized to form an oxide adlayer on the nanocrystal surfaces as the applied potential exceeded certain threshold values. Because the undercoordinated surface atoms at the surface defects, domain boundaries, and locally curved surface sites were oxidized more easily than their counterparts on the atomically flat terraces, the oxidation potential downshifted as the surface atomic coordination number decreased.^46–49^ As shown in Figure 3D, two characteristic oxidation peaks during the anodic potential sweep were centered at 1.00 and 1.18 V *vs.* saturated calomel electrode (SCE), signifying the electrochemical oxidation of undercoordinated Au surface atoms. In contrast, the close-packed surface atoms on the terraces or crystallographic facets, typically the {111} and {100} facets, were oxidized in a more positive potential range above ∼ 1.3 V (vs. SCE). The CV results clearly revealed that the surfaces of CNM-Au8 had a high abundance of undercoordinated surface atoms. During the cathodic potential sweep, a sharp reduction peak emerged within the potential window from ∼1.00 to ∼ 0.85 V (vs. SCE), which was the electrochemical signature of the oxide adlayer stripping. Assuming a specific charge of 450 μC cm^−2^,^50^ the mass-specific electrochemically active surface area (ECSA) of CNM-Au8 nanocrystals was estimated to be ∼23.1 ± 1.1 m^2^ g^−1^, in fairly good agreement with the theoretically predicted mass-specific surface area of an Au nanosphere with a diameter of 11.3 nm (27.6 m^2^ g^−1^).

**Figure 3.**
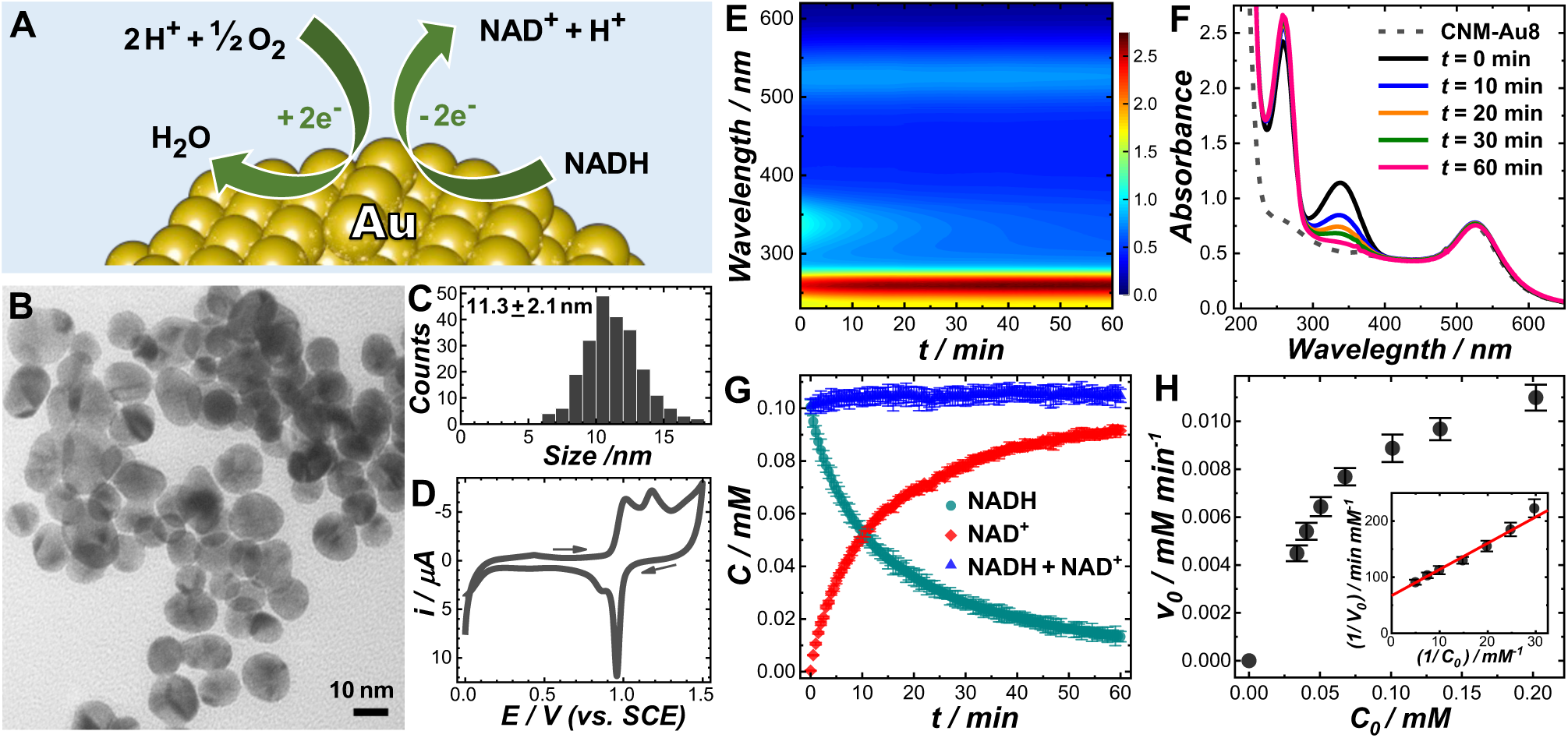
(A) Schematic illustration of Au-catalyzed NADH oxidation. (B) Transmission electron microscopy (TEM) image and (C) particle size distribution of CNM-Au8. (D) Cyclic voltammetry curve collected from a CNM-Au8-loaded work electrode in an N_2_-purged aqueous electrolyte containing 0.5 M H_2_SO_4_ at room temperature. The potential sweep rate was 5.0 mV s^-1^ and the arrows showed the direction of potential sweep. (E) Temporal evolution of UV-vis absorption spectra and (F) snapshot UV-vis spectra at various reaction times during the CNM-Au8-catalyzed NADH oxidation at room temperature in a neutral environment (pH of 7). The mass concentration of Au catalysts was 37.7 mg L^-1^ and the initial concentration of NADH was 0.10 mM. (G) Temporal evolution of NADH concentration, NAD^+^ concentration and the total concentration of NADH and NAD^+^ during the CNM-Au8-catalyzed NADH oxidation reaction at a pH of 7. (H) Relationship between the initial reaction velocity and the initial concentrations of NADH. The Lineweaver-Burk plot is shown in the inset. The curve fitting result is shown as a solid red line in the inset of panel H.

We used UV-Vis absorption spectroscopy as an *in situ* spectroscopic tool to monitor the progress of Au-catalyzed NADH oxidation in real time. NADH has two characteristic absorption peaks centered at the wavelengths of 340 and 260 nm, respectively. The peak at 340 nm is the spectral signature of the n–π* transition of the dihydronicotinamide unit, while the absorption peak at 260 nm corresponds to the π–π* transition of the adenine ring.^51^ The molar absorption coefficients of NADH at 340 and 260 nm are 6220 and 14100 M^-1^cm^-1^, respectively.^52^ The oxidized product, NAD^+^, also strongly absorbs light at 260 nm, with a molar absorption coefficient (16900 M^-1^cm^-1^) even higher than that of NADH. Figure 3E shows the temporal evolution of UV-vis absorption spectra during CNM-Au8-catalyzed aerobic oxidation of NADH in a neutral aqueous medium (pH of 7) at room temperature (22 °C). Several snapshot spectra captured at various reaction times are highlighted in Figure 3F. The initial molarity of NADH was 0.10 mM and the mass concentration of CNM-Au8 nanocatalysts was kept at 37.7 mg L^-1^ (0.191 mM Au atoms). The light extinction peak centered at ∼528 nm, which was spectral feature of the localized plasmon resonances sustained by CNM-Au8 nanocrystals, remained essentially unshifted with negligible intensity changes (Figure 3E and 3F), indicating that colloidal Au nanocatalysts remained well-dispersed without forming aggregates during the reaction. As the catalytic reaction proceeded, the absorption intensity at 340 nm progressively decreased while the absorption peak at 260 nm became increasingly more intense. Based on the Lambert-Beer’s law and the molar absorption coefficients of NADH and NAD^+^, we were able to further convert the experimentally measured absorbance to the molar concentrations of NADH and NAD^+^. As shown in Figure 3G, the decrease in NADH concentration, *C*(NADH), was accompanied by the increase in NAD^+^ concentration, *C*(NAD^+^), with the sum of *C*(NADH) + *C*(NAD^+^) kept almost unchanged throughout the entire reaction process, indicating that neither long-lived intermediates nor side products were observed in the UV-Vis absorption spectroscopic results. One of the key parameters for analyzing the kinetic results was the initial reaction velocity, *v_0_*, whose value could be extracted by fitting the temporal evolution of *C*(NADH) at the initial stage of the reactions with a linear function. When keeping the mass concentration of the CNM-Au8 nanocatalysts fixed at 37.7 mg L^-1^ and the pH fixed at 7, the relationship between *v_0_* and the initial NADH concentration, *C_0_*, could be well-described by the steady-state Michaelis-Menten kinetic model (Figure 3H), indicating that CNM-Au8 essentially functioned as a nanozyme (enzyme-mimicking nanocatalyst) while NADH served as a substrate to the nanozyme. A linear relationship between the reciprocal of *v_0_* and the reciprocal of *C_0_*was clearly observed (inset of Figure 3H), which allowed us to obtain the values of the Michaelis constant, *K_m_*, and the maximal velocity, *v_max_*, by fitting the experimental results with the Lineweaver-Burk equation:

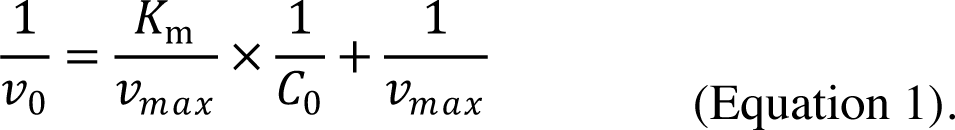

At a fixed mass concentration of CNM-Au8 and initial concentration of NADH, the reaction rate increased significantly as the reaction medium became more acidic (Figure 4A) because the catalytic NADH oxidation reactions involved protons and the affinity of reactants to the Au nanozyme surfaces was pH-dependent. Although the reaction rates changed significantly upon variation of pH, the kinetic features of the reactions could always be well-described using the Michaelis-Menten model over a broad range of pHs from 3 to 9 (Figure 4B and 4C). As the pH progressively increased, the values of *v_max_* extracted from least-squares curve fitting decreased monotonically until reaching a plateaued value around 0.015 mM min^-1^ at pHs above 7 (Figure 4D). The values of *K_m_* remained close to 0.1 mM in acidic environments at pHs below 7, whereas a significant increase of *K_m_* was observed upon further elevation of pH in the alkaline environment (Figure 4E). The pH-dependence of *v_max_*and *K_m_* shown in Figure 4D and 4E indicated that increase of pH values not only led to decrease of the turnover number, *k_cat_*, which was defined as *v_max_* normalized against the nanozyme concentration, *C_nanozyme_*, but also weakened the interactions between the NADH substrate and the Au nanozyme. We further calculated the catalytic efficiency, *k_cat_*/*K_m_*, at various pHs using the following equation:

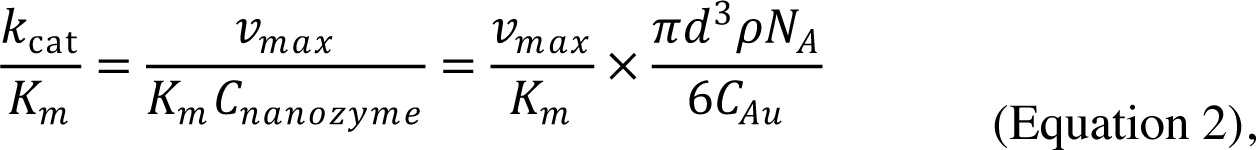

**Figure 4.**
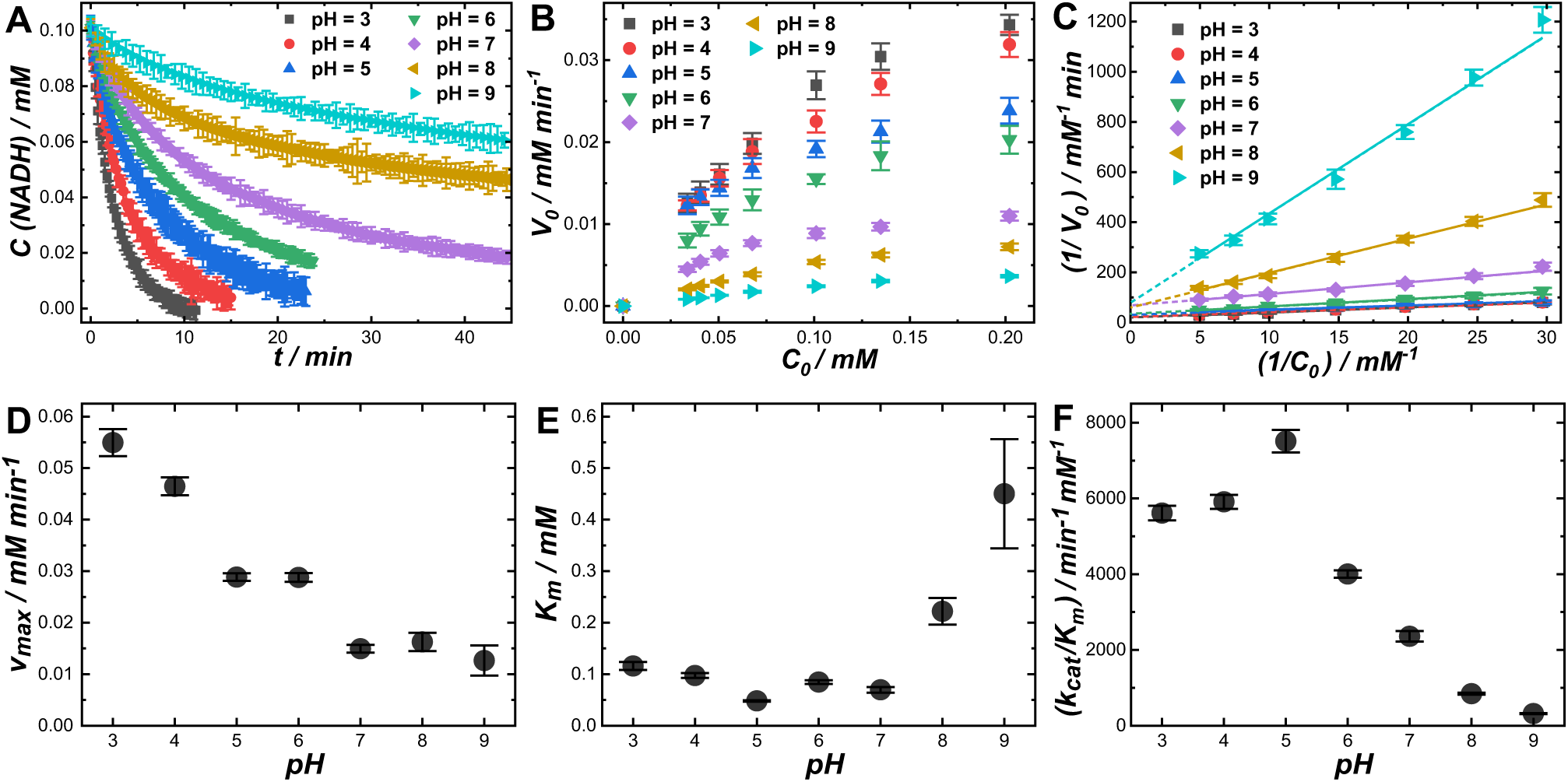
(A) Temporal evolutions of NADH concentration during CNM-Au8-catalyzed NADH oxidation reactions at various pHs at room temperature. The mass concentration of Au catalysts was 37.7 mg L^-1^ and the initial concentration of NADH was 0.10 mM. (B) Relationships between *v*_0_ *and C*_0_ at various pHs. (C) Lineweaver-Burk plots at various pHs. The reactions occurred at room temperature. pH-dependence of (D) *v_max_*, (E) *K_m_*, and (F) *k_cat_/K_m_*.

in which *d* is the average diameter of CNM-Au8 nanocrystals (11.3 nm), 𝜌 is the mass density of Au (19.3 g cm^-3^), *N_A_* is the Avogadro′s number (6.02 ×10^23^ mol^-1^), and *C_Au_* is the mass concentration of Au in the reactant mixtures (37.7 mg L^-1^). CNM-Au8 appeared catalytically more efficient in acidic environments than in alkaline environments, reaching the maximal catalytic efficiencies around pH of 5 (Figure 4F). While there is typically one active site on each natural enzyme molecule, each CNM-Au8 nanocrystals may have multiple active sites on its surface. To assess the intrinsic catalytic efficiency at a single active site, the calculated values of *k_cat_*/*K_m_* need to be further normalized against the number of active sites on each CNM-Au8 nanocrystal, which still remains unknown at this point.

The pH-dependent catalytic behaviors of CNM-Au8 were further studied through electrochemical measurements. Figure S3 in the Supporting Information shows the CV curves collected from glassy carbon electrodes (GCEs) and GCEs loaded with CNM-Au8 in electrolytes containing 1 mM NADH at various pHs. In acidic electrolytes, both the onset potential, *E_onset_*, and the peak potential, *E_p_*, for the NADH oxidation were shifted negatively when CNM-Au8 was loaded onto the working electrodes, suggesting that CNM-Au8 was capable of catalyzing the NADH oxidation. Because of the involvement of protons, *E_p_* became more positive as pH increased. In acidic electrolytes, *E_p_* exhibited lower values on CNM-Au8-loased GCEs than on GCEs, whereas such differences in *E_p_* values became negligible in alkaline environments at pHs above 7. The results of the electrochemical measurements further confirmed that the nanocrystals in CNM-Au8 were catalytically more active toward NADH oxidation in acidic environments than in alkaline environments. Regardless of pH, the electrochemical oxidation of NADH on GCEs and CNM-Au8 exhibited diffusion-controlled kinetic features, as the anodic peak current, *i_p_*, appeared proportional to the square-root of the potential sweep rate, *v*^1/2^, in the linear sweep voltammetry (LSV) results (Figure S4 in the Supporting Information).

The CNM-Au8-catalyzed NADH oxidation reactions could be kinetically modulated not only by adjusting pH but also by varying the concentration of dissolved molecular O_2_ in the aqueous medium (Figure S5 in the Supporting Information). When purging the catalyst-reactants mixtures by displacing ambient air with O_2_, the reactions became kinetically faster than those occurring under ambient air. In contrast, purging the catalyst-reactants mixtures with N_2_ led to slower reaction rates due to decreased concentrations of dissolved O_2_. These observations strongly indicated that both NADH and O_2_ were substrate molecules to the Au nanozymes. An enzymatic reaction involving two different substrate molecules may occur through either a ping-pong mechanism, in which the reaction proceeds with the release of one product prior to the association of the other substrate, or a sequential mechanism, which requires the co-adsorption of both substrates to the enzyme before the product molecules are generated and released from the enzyme. These two fundamentally distinct mechanisms could be distinguished from each other by inspecting the Lineweaver-Burk plots at various concentrations of dissolved O_2_ while keeping the initial concentration of NADH fixed. For a ping-pong mechanism, such a plot will yield a series of straight lines that are parallel to each other because the slopes are independent of O_2_ concentrations. However, the Lineweaver-Burk plots for the sequential mechanism yield a family of unparallel straight lines that intersect to the negative side of the (1/*v*_0_) axis. The kinetic results shown in Figure S5 in the Supporting Information revealed that the CNM-Au8-catalyzed NADH oxidation involved the formation of a nanozyme-NADH-O_2_ ternary complex, following the sequential mechanism rather than the Ping Pong mechanism.

The overall catalytic activities of CNM-Au8 were intimately tied to the atomic level surface structures of the nanocrystals. We observed a gradual decay in the catalytic activities of CNM-Au8 over multiple cycles of reactions at a pH of 3 (Figure S6 in the Supporting Information). In each reaction cycle, we monitored the temporal evolution of NADH at a fixed mass concentration of CNM-Au8 of 37.7 mg L^-1^ and initial NADH concentration of 0.10 mM. After NADH was completely converted into NAD^+^, a new cycle of catalytic reaction was initiated upon introduction of additional NADH, which brought the NADH concentration back to 0.10 mM. The activity decay over multiple reaction cycles was a ubiquitous phenomenon observed over a broad range of pHs (Figure S7 in the Supporting Information), which was found to be related to the atomic level surface restructuring of nanocrystals of CNM-Au8 during the catalytic reactions. Although the CNM-Au8 nanocrystals recycled after 5 cycles of reactions at pH of 3 exhibited no observable changes in particle sizes or morphologies in the TEM images in comparison to the as-synthesized CNM-Au8, the results of CV-based surface oxide stripping measurements showed that the fraction of undercoordinated Au surface atoms decreased considerably after 5 reaction cycles, while the mass-specific ECSA was well-preserved (Figure S8 in the Supporting Information). The above-mentioned observations strongly suggested that the undercoordinated surface atoms served as the primary active sites for the catalytic NADH oxidation reactions. Due to the absence of organic capping ligands on the CNM-Au8 nanocrystal surfaces, these active sites were easily accessible to reactant molecules. In Figure S9 in the Supporting Information, we further compared the catalytic performances of CNM-Au8 to those of Au quasi-spherical nanoparticles (QSNPs, 11.1 ± 0.4 nm in diameter) and surface-roughened nanoparticles (SRNPs, 152 ± 4.8 nm in diameter), whose surfaces were coated with a bilayer of capping surfactants, cetyltrimethylammonium chloride (CTAC). Although CNM-Au8 and the CTAC-capped Au QSNPs exhibited similar particle sizes and morphologies, the surface capping of CTAC led to remarkably decreased catalytic activity. As shown in detail in our previous publications,^53–56^ the Au SRNPs had high abundance of undercoordinated surface atoms, well-mimicking the locally curved surface structures of sub-10 nm nanoparticles. However, due to presence of surface-capping CTAC molecules and significantly larger particle sizes, the Au SRNPs exhibited a mass-specific catalytic activity drastically lower than that of CNM-Au8. Therefore, highly abundant undercoordinated surface atoms and organic ligand-free clean surfaces of CNM-Au8 were both key contributory factors leading to the high catalytic activities toward NADH oxidation.

### Kinetic Enhancements Under Light Illumination

Au nanoparticles exhibit intriguing optical properties that are dominated by the collective oscillation of free electrons, also known as plasmons. The photoexcitation and decay of plasmons give rise to a cascade of photophysical processes, including enhancements of local fields on nanoparticle surfaces, generation of non-equilibrium hot charge carriers, and photothermal transduction, all of which may play crucial roles in driving or enhancing photocatalytic transformations of molecular adsorbates on the metal nanocrystal surfaces.^57–70^ Because the particle size of CNM-Au8 (below 20 nm) is well within the quasi-static limit, CNM-Au8 nanocrystals behave as strong visible light absorbers upon resonant excitation of their plasmons with negligibly small contribution of scattering to the overall light extinction. Although limited local-field enhancements are achievable on the surfaces of small Au nanoparticles, CNM-Au8 is expected to be more efficient in producing hot charge carriers than their counterparts with larger particle sizes, which is beneficial for plasmonic hot carrier-driven photocatalytic reactions. Au nanoparticles also strongly absorb light on the higher-energy side of their plasmon resonances due to interband electronic transitions from the valence band (5d) to the conduction band (6sp).^71–73^ When the excitation photon energy exceeds the interband energy threshold of Au (energy above ∼2.4 eV, wavelength below ∼516 nm), hot electrons and holes can also be generated upon excitation of d-to-sp interband transitions. While the interband hot carriers are typically considered to have lower kinetic energies as those of the intraband plasmonic hot carriers, they can also be effectively harnessed to drive photocatalytic reactions, achieving even higher photocatalytic efficiencies than those of plasmon-driven reactions in certain cases.^71, 72^ The internal decay of the plasmonic and interband hot carriers within a metallic nanoparticle results in local heating of the nanoparticle itself, followed by heat dissipation to its surrounding medium, providing a unique pathway to deposit thermal energies into the vibrational modes of molecular adsorbates and accelerate molecular transformations on the nanoparticulate photocatalyst surfaces.^70, 74^ In this work, we used both the hot carriers and photothermal heating resulting from interband and plasmonic excitations as unique leverages to kinetically maneuver the CNM-Au8-catalyzed NADH oxidation reactions at a remarkable level of precision.

We chose continuous wave (CW) lasers with emission lines at 445 and 520 nm as the light sources to excite the interband transitions and plasmon resonances of the nanocrystals in CNM-Au8, respectively, while a CW excitation laser with an emission line at 785 nm was used for the control experiments under off-resonance conditions (Figure 5A). The mass concentration of CNM-Au8 nanocatalysts was fixed at 37.7 mg L^−1^ with a total volume of the catalyst-reactant mixtures kept at 2.0 mL. The excitation lasers were collimated with a 4 mm × 4 mm, square-shaped illumination cross-section and incident vertically from the top surface into 2 mL of catalyst-reactants mixtures in a 1.0 cm × 1.0 cm × 4.5 cm quartz cuvette. The power of the excitation lasers, *Pex*, was adjusted in the range of 0-1.0 W. The catalyst-reactants mixtures were kept under constant magnetic stir (800 rpm) to facilitate heat dissipation such that thermal equilibria were rapidly established between the nanocrystals and the bulk solution. To eliminate the temperature elevation caused by photothermal transduction, a circulating water bath was used to maintain an isothermal reaction condition by controlling the bulk solution-phase temperature at 22 ± 0.5 °C. As shown in Figure 5B and 5C, the CNM-Au8-catalyzed reactions were significantly accelerated under illuminations by the 445 nm and 520 nm lasers, and increasing the illumination power led to increased reaction rates. However, at an excitation wavelength (*λex*) of 785 nm, the reaction rates under photo-illumination and in dark were observed to be almost the same (Figure 5D) because the excitation photons were off-resonant with the plasmons and energetically insufficient for the excitations of the interband transitions. We fitted the experimentally measured *Pex*-dependence of reaction rates at the initial stage of the reactions with following power function (Figure 5E):

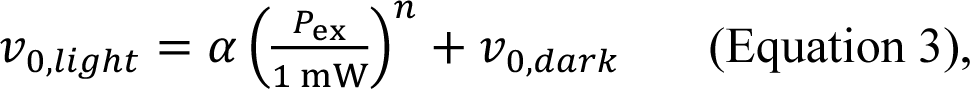

**Figure 5.**
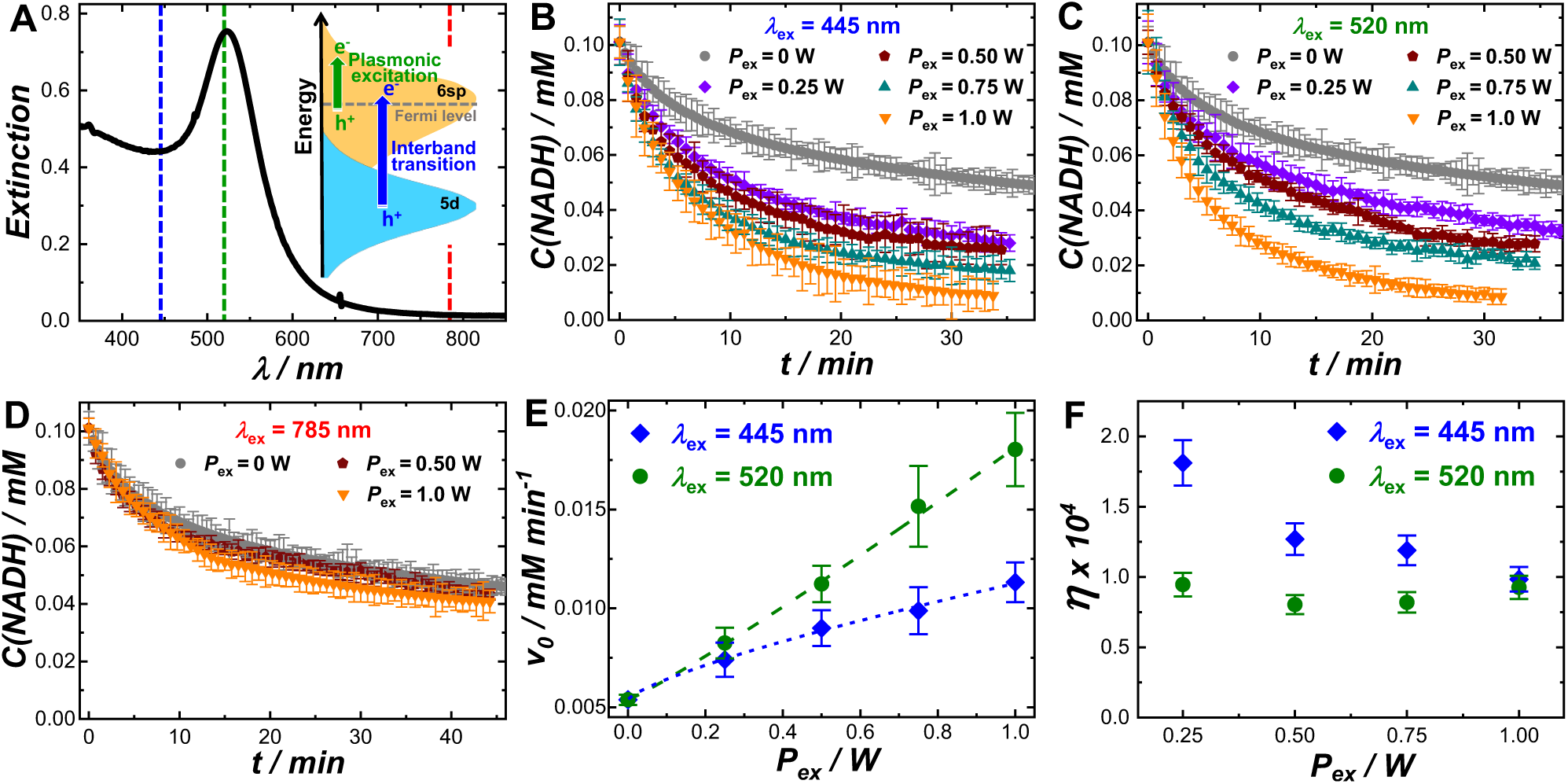
(A) Light extinction spectrum of colloidal CNM-Au8. The vertical dash lines indicate the wavelengths of the excitation lasers used in this work. The inset schematically illustrates the photoexcitation of hot electrons and holes upon excitations of the interband transitions and the intraband plasmon resonances. Temporal evolutions of NADH concentration during CNM-Au8-catalyzed NADH oxidation reactions under photo-illumination at *λex*s of (B) 445, (C) 520, and (D) 785 nm and various *Pex*s. The reactions occurred at a pH of 8 under isothermal conditions with temperature controlled at 22 ± 0.5 °C. The mass concentration of Au catalysts was 37.7 mg L^-1^ and the initial concentration of NADH was 0.10 mM. *Pex*-dependence of (E) *v0* and (F) *η* at *λex*s of 445 and 520 nm. The experimentally measured *Pex*-dependence of *v0* was fitted with Eqn.5 and the curve fitting results were shown as dash curves in panel E.

in which 𝑣_0,*light*_ and 𝑣_0,*dark*_ refer to the initial reaction velocities under photo-illumination and in dark, respectively. *α* is a fractional coefficient and *n* is an exponent. 𝑣_0,*light*_ was observed to increase almost linearly with *Pex* when the plasmon resonances were excited at *λex* of 520 nm, and the *n* value obtained from least squares curve fitting (1.09 ± 0.04) was very close to 1. In contrast, a sublinear *Pex*-dependence of reaction rates was clearly observed with an *n* value of 0.75 ± 0.05 at *λex* of 445 nm when the interband transitions were optically excited. Such sublinear *Pex*-dependence of reaction rates is also a common kinetic feature of exciton-driven photocatalysis on semiconductors. ^62, 75^

We further compared the apparent quantum efficiencies, *η*, for the reaction kinetic enhancements under the intraband plasmonic (*λex* = 520 nm) and interband (*λex* = 445 nm) excitations (Figure 5F). Here, *η* was defined as the ratio between the number of electrons harnessed for the reactions and the number of photons absorbed by the CNM-Au8 nanocrystals over a reaction time of 360 s. *η* was calculated using the following equation:

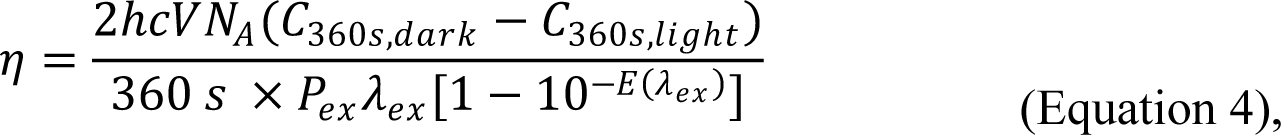

where *h* is Planck’s constant (6.626 × 10^−34^ m^2^ kg s^−1^), *c* is the speed of light (3.0 × 10^8^ m s^−1^), *V* is the total volume of the catalyst-reactants mixtures (2.0 × 10^−3^ L), *NA* is Avogadro’s number (6.022 × 10^23^ mol^−1^), *C*360s,light and *C*360,dark are the concentrations of NADH at a reaction time of 360 s under light illumination and in dark, respectively. *E*(*λex*) is the optical extinction at the excitation wavelength, *λex*, with an optical path length of 2 cm. When *Pex* was varied within the sub-mW range, *η* decreased as *Pex* increased at *λex* of 445 nm, whereas *η* appeared independent of *Pex* at *λex* of 520 nm. The linear power-dependence of reaction rates and power-independent quantum efficiencies have been considered as kinetic signatures of plasmonic hot carrier-driven reactions under isothermal conditions.^62, 75, 76^ For photocatalytic reactions driven by interband excitations, the thermalization or recombination of the interband hot carriers becomes prevalent in the high-power regime, leading to sublinear power dependence of reaction rates and decreased quantum efficiencies at higher excitation powers.^71^

In addition to the plasmonic and interband hot carriers, the photothermal heating effects could also be harnessed to kinetically boost the CNM-Au8-catalyzed NADH oxidation reactions. We systematically studied the kinetics of the reactions taking place in ambient air under continuous laser illuminations without using the temperature-controlled circulating water bath. We consistently observed that the reactions became faster under ambient air at *λex*s of both 445 and 520 nm when the circulating water bath was removed (Figure 6A and 6B and Figure S10 in the Supporting Information). Because the thermal conductivity of air was substantially lower than that of water, the dissipation of the heat generated from photothermal transduction became significantly slower, which resulted in elevation of the temperatures in the catalyst-reactants system. We monitored the temperature evolutions of colloidal CNM-Au8 (37.7 mg L^-1^) and pure water under continuous laser illuminations at various *P_ex_*s until the temperature reached plateaued values, *T_eq_*, at the thermal equilibria, followed by natural cooling of the samples down to room temperature in dark (Figure 6C and 6D). Considerably higher temperatures were reached at the thermal equilibria in colloidal CNM-Au8 than in water because of photothermal transduction resulting from the interband and plasmonic excitations of the colloidal Au nanoparticles. The temperature elevation caused by photothermal transduction of CNM-Au8 (*T_eq_* of colloidal suspensions of CNM-Au8 subtracted by *T_eq_* of pure water, *T_eq,H_2_O_*) was found to be linearly dependent on *P_ex_* for both the interband and plasmonic excitations (Figure 6E).

**Figure 6.**
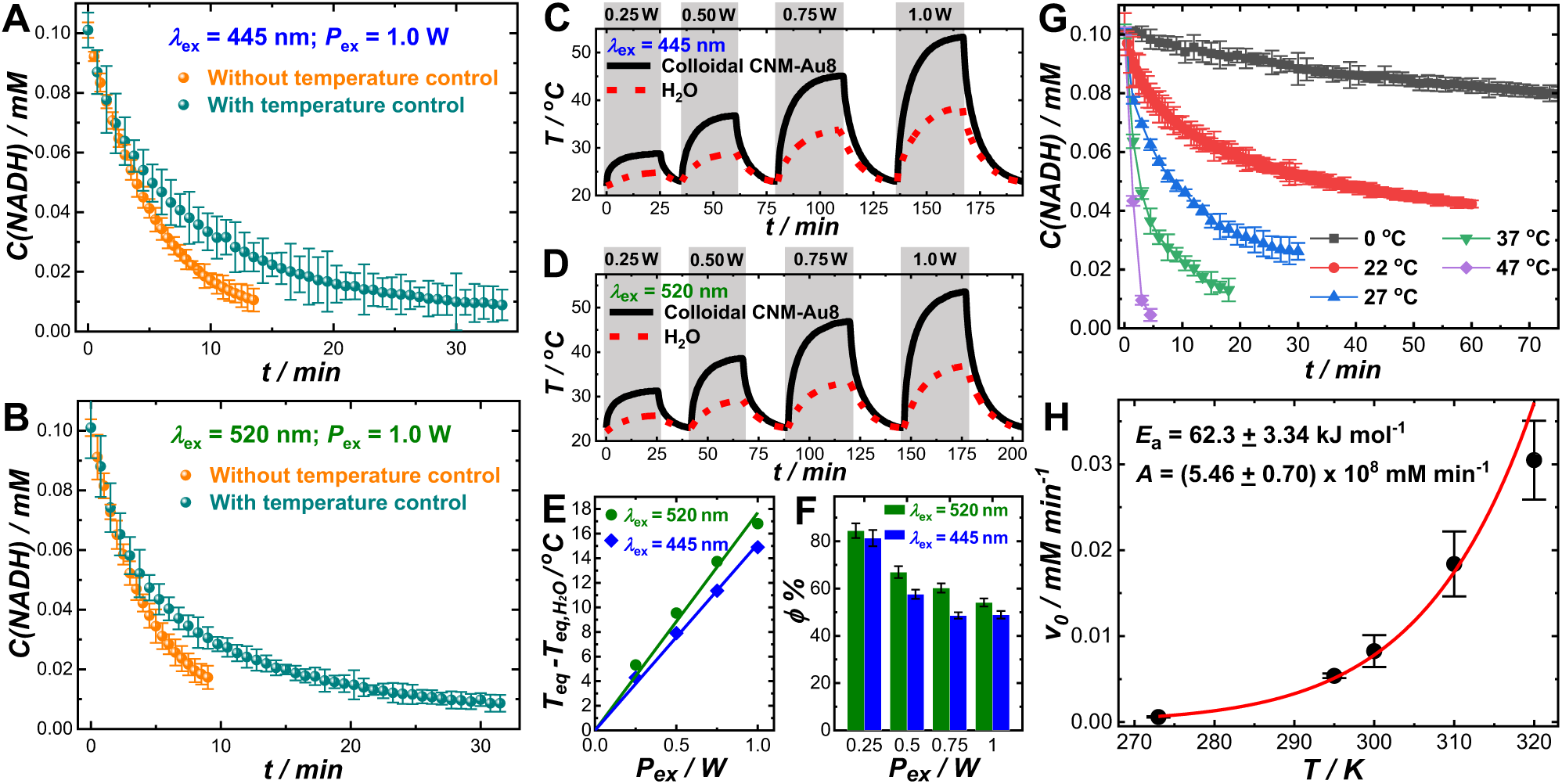
Temporal evolutions of NADH concentration during CNM-Au8-catalyzed NADH oxidation reactions under photo-illumination at a *Pex* of 1.0 W and *λex*s of (A) 445 and (B) with and without temperature control. Temporal evolutions of the bulk temperatures of colloidal CNM-Au8 (37.7 mg L^-1^) and water under multiple cycles of photo-illumination by the (C) 445 and (D) 520 nm lasers at various *P_ex_*s and natural cooling in dark under ambient air. *P_ex_*-dependence of (E) *T_eq_* - *T_eq,H_2_O_* and (F) *ϕ*. (G) Temporal evolutions of NADH concentration during CNM-Au8-catalyzed NADH oxidation reactions in dark at various reaction temperatures. (H) Relationship between *v0* and reaction temperature at a pH of 8. The experimental results were fitted with the Arrhenius equation. In all reactions, the mass concentration of Au catalysts was kept at 37.7 mg L^-1^ and the initial concentration of NADH was 0.10 mM.

We calculated the photothermal transduction efficiency, *ϕ*, based on a widely used thermophysical model.^77–79^ The values of the thermal equilibrium time constant, *τs*, were obtained by fitting the temporal evolution of -ln(*θ*) during natural cooling in dark with a linear function (Figure S11 in the Supporting Information):

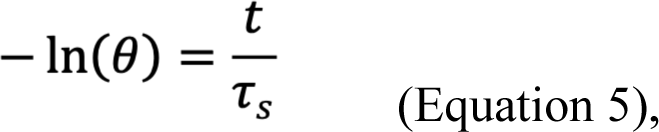

where 𝜃 refers to the driving force toward thermal equilibrium, which is defined as

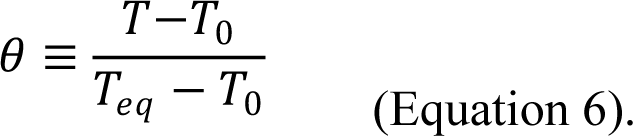

Here, *T_0_* is the ambient temperature (22 °C in our case). At the thermal equilibrium, the heat generation and dissipation occur at the same rate, and *ϕ* can be calculated using the following equation:

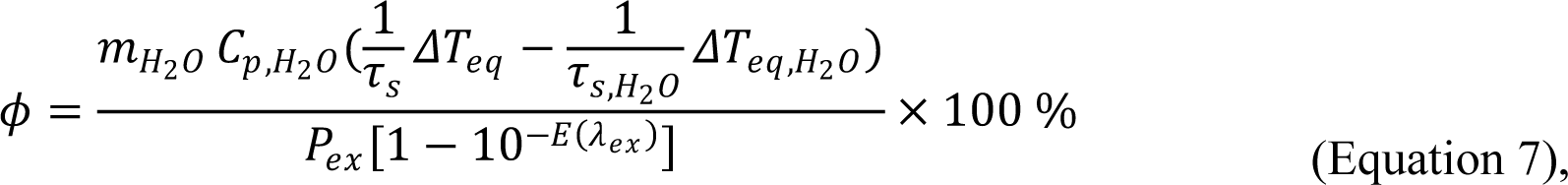

in which *mwater* and *Cp,water* are the mass and constant-pressure heat capacity of water (*mwater* = 2.0 g and *Cp,water* = 4.184 J K^-1^ g^-1^), respectively. *ΔTeq* and *ΔTeq,H2O* represent temperature elevation of colloidal suspensions of CNM-Au8 and pure water, respectively, at the thermal equilibrium. *τs,water* refers to the *τs* of water in the absence of CNM-Au8 nanocrystals. As shown in Figure 6F, *ϕ* values of plasmonic excitations at *λex* of 520 nm were slightly higher (within 10 %) than those of interband excitations at *λex* of 445 nm at identical *Pex*s, with higher *ϕ* values observed at lower *Pex*s at each *λex*. Comparison of the reaction rates under photo-illuminations with and without temperature control, together with the photothermal results, strongly indicated that photothermal heating kinetically boosted the CNM-Au8-catalyzed NADH oxidation reactions.

To further confirm the effects of temperature elevation on the kinetic enhancements, we studied the kinetics of dark reactions without laser illumination at various constant temperatures by immersing the catalyst-reactants mixtures in a temperature-controlled water bath. As shown in Figure 6G, increase of the reaction temperature led to significant acceleration of the reactions at the pH of 8. At a fixed initial concentration of NADH and a fixed catalyst mass concentration, 𝑣_0_ becomes directly proportional to the rate constant of the rate-limiting step, which is the *kcat* of a Michaelis-Menten enzymatic reaction. Therefore, the relationship between *v*0 and the temperature, *T*, could be well-described by the Arrhenius equation:

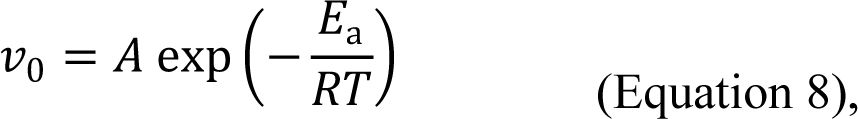

in which *Ea* is the apparent activation energy of the rate-limiting step, *R* is the molar gas constant (8.314 J mol^-1^ K^-1^), and *A* is a pre-exponential factor. By fitting the experimental results with the Arrhenius equation, the value of *Ea* at pH of 8 was determined to be 62.3 ± 3.34 KJ mol^-1^. At pH of 5, the Arrhenius relationship between *v*0 and *T* was still held (Figure S12 in the Supporting Information). Switching the pH from 8 to 5 resulted in a decrease of *Ea* by ∼ 12.1 KJ mol^-1^, fully in line with the pH-dependent catalytic behaviors of CNM-Au8 observed in this work.

## CONCLUSIONS

We have found that a pharmaceutical grade suspension of electrochemically synthesized Au nanocrystals, referred to as CNM-Au8, may function as a NADH dehydrogenase-mimicking nanozyme that regulates the bioenergetic metabolism and thereby effectively protects the central nervous system cells in a manner agnostic to neuron type. The neuroprotective function of CNM-Au8 includes aerobic oxidation of NADH, catalyzed by the CNM-Au8 nanocrystals. This oxidation reaction by CNM-Au8 nanocrystals obeys the Michaelis-Menten kinetic model. The high abundance of undercoordinated surface atoms and the organic ligand-free clean surfaces of the nanocrystals in CNM-Au8 have been found to be two critical factors leading to the remarkable catalytic activities of CNM-Au8 toward NADH oxidation. The insights gained from this work provide important guiding principles for rational design of noble metal-based nanozymes with bioenergetic metabolism-regulating and neuroprotective functions. We have further demonstrated that photoexcited charge carriers and photothermal transduction, which are derived from optical excitations of either the plasmonic electron oscillations or the interband electronic transitions in CNM-Au8, can be judiciously harnessed to further enhance the reaction kinetics under visible light illumination in a highly controllable manner. By deliberately tailoring the sizes and shapes of clean-surfaced gold nanocrystals, it becomes possible to tune the plasmon resonance frequencies over a broad spectral range across the visible spectral region into the near-infrared water window, in which tissues and blood become transparent. Using the deep-penetrating near-infrared light to fine-regulate the bioenergetic metabolism in living cells opens up a new avenue for further improving the efficacy of Au nanoparticle-based therapeutic agents for neurodegenerative diseases.

## METHODS

The chemicals and reagents were used as received without further purification (see a detailed list in the Supporting Information). Ultrapure water with a resistivity of 18.2 MΩ (Barnstead EasyPure II 7138) was used for all experiments. CNM-Au8 was synthesized using a previously published and patented electrochemical method.^35, 37^ The CNM-Au8 nanoparticles were characterized by TEM, light extinction spectroscopy, and electrochemical measurements, as detailed in the Supporting Information. CTAC-capped Au QSNPs^80^ and SRNPs^53–56^ were synthesized following previously published protocols (see details in the Supporting Information). The protocols for the *in vitro* neuroprotection assays were described in detail in the Supporting Information.

The progress of NADH oxidation reactions under various conditions was tracked in real time based on the temporal evolution of NADH concentrations using an Agilent 8453 UV-Vis-NIR spectrophotometer. The catalytic reactions were carried out in a 1 cm × 1 cm × 4.5 cm quartz cuvette containing various initial concentrations of NADH in citrate buffers at various pHs. The total volume of the catalyst-reactants mixtures was kept at 2.0 mL. The mass concentration of CNM-Au8 was 37.7 mg L^-1^. The reactions occurred under an isothermal condition using a temperature-controlled water bath. CW lasers purchased from Lasever Inc. (Ningbo, Zhejiang, China) with the tunable power output in the range of 0-3.0 W and emission wavelengths at 445nm (model no. LSR445SD), 520nm (model no. LSR520CPD), and 785 nm (model no. LSR785NL) were used as the light sources for the photo-illuminated reactions. The excitation lasers were collimated with a 4 mm × 4 mm, square-shaped illumination cross-section and incident vertically from the top surface into 2 mL of catalyst-reactants mixtures. The catalyst-reactants mixtures were kept under constant magnetic stir (800 rpm) to facilitate heat dissipation. For photo-illuminated reactions under isothermal conditions, a circulating water bath was used to control the bulk solution-phase temperature at 22 ± 0.5 °C. The temperature evolution during photo-illumination and natural cooling was monitored using a digital thermocouple (Therma Waterproof Thermometer for Type K Thermocouples, Priggen Special Electronic).

## ASSOCIATED CONTENT

### Supporting Information

Additional experimental details: chemicals and reagents, nanoparticle synthesis, nanoparticle characterizations, and detailed protocols for the *in vitro* neuroprotective assays. Additional figures as noted in the maintext: results of neuroprotective assays, electrochemical results, kinetic results, and photothermal results. This material is available free of charge via the Internet at http://pubs.acs.org.

## AUTHOR INFORMATION

### Corresponding Authors

- Karen S. Ho - Clene Nanomedicine, Inc., Salt Lake City, Utah 84117, United States; orcid.org/0000-0003-3217-354X
- Hui Wang − Department of Chemistry and Biochemistry, University of South Carolina, Columbia, South Carolina 29208; Email; orcid.org/0000-0002-1874-5137

### Authors

Zixin Wang − Department of Chemistry and Biochemistry, University of South Carolina, Columbia, South Carolina 29208; orcid.org/0000-0002-3604-860X.

Alexandre Henriques - Neuro-Sys, 410 D60, 13120 Gardanne, France Laura Rouvière - Neuro-Sys, 410 D60, 13120 Gardanne, France Noëlle Callizot - Neuro-Sys, 410 D60, 13120 Gardanne, France

Lin Tan - Department of Chemistry and Biochemistry, University of South Carolina, Columbia, South Carolina 29208

Michael T. Hotchkin - Clene Nanomedicine, Inc., Salt Lake City, Utah 84117, United States

Mark G. Mortenson - Clene Nanomedicine, Inc., Salt Lake City, Utah 84117, United States and Clene Nanomedicine, Inc., North East, Maryland 21901, United States

Rodrigue Rossignol - Cellomet, Functional Genomics Center (CGFB), 146 rue Léo Saignat, 33000 Bordeaux, France

### Author Contributions

K.S.H., M.T.H., and H.W. designed the project. A.H., L.R., and N.C. performed the *in vitro* experiments on primary neuronal-glial co-cultures. R.R. conducted the ATP assay. Z.W. characterized CNM-Au8 and its reaction kinetics. L.T. compared the catalytic performance of CNM-Au8 to other Au nanoparticle samples. M.G.M. originated the methods for the electrochemical synthesis of CNM-Au8. All authors discussed the results and participated in data analysis. K.S.H. and H.W. wrote the paper with contributions from all authors. All authors have given approval to the final version of the manuscript.

### Notes

Z.W., L.T., and H.W. declare no competing financial interest. K.S.H., M.T.H., and M.G.M. are full time employees of Clene Nanomedicine, Inc. and own shares in the company. A.H., L.R., and N.C. are full-time employees of Neuro-Sys, a private contract research organization. R.R. is the co-founding CEO and CSO of CELLOMET, a private contract research organization.

## ACKNOWLEDGMENT

Clene Nanomedicine, Inc. funded this research. The authors thank Random42 and Bjorn Pendleton (Pendleton Creative, LLC) for the TOC artwork. This work used the Hitachi HT7800 transmission electron microscope in the Electron Microscopy Center at University of South Carolina, which was purchased using funds provided by an NSF EPSCoR RII Track-I Award (OIA-1655740).

## TOC Graphic

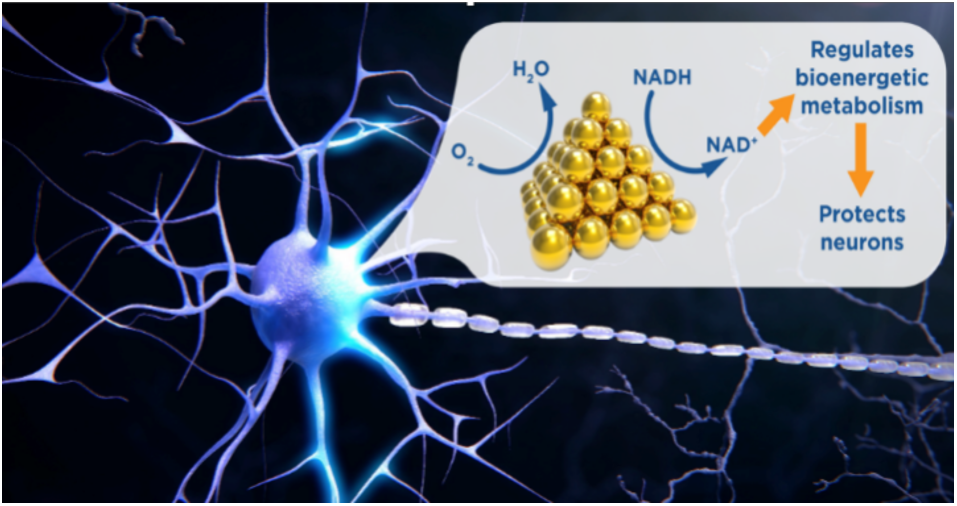

