## Supplemental Information for "A Mechanism Underpinning the Bioenergetic Metabolism-Regulating Function of Gold Nanocatalysts"

**± Clene Nanomedicine, Inc., North East, Maryland 21901, United States**

[#] Cellomet, Functional Genomics Center (CGFB), 146 rue Léo Saignat, 33000 Bordeaux, France

* Corresponding Authors:

 (K.S. Ho)

 (H.Wang)

¶ These authors contributed equally.

### S1. Additional Experimental Details

#### S1.1. Nanoparticle Synthesis

**Chemicals and Materials.** CNM-Au8® (500 mg $L^{-1}$) and the vehicle solution (6.5 mM $NaHCO_3$ in pharmaceutical-grade water) were provided by Clene Nanomedicine, Inc (CNM-Au8® is a federally-regulated trademark of Clene Nanomedicine, Inc). Gold(III) chloride trihydrate ($HAuCl_4 \cdot 3H_2O$, ACS grade) was purchased from J.T. Baker. Sulfuric acid ($H_2SO_4$, 98 %), sodium hydroxide (NaOH), sodium citrate dihydrate, and ethanol (200 proof) were purchased from Fisher Scientific. Sodium borohydride ($NaBH_4$, 99.99 % trace metals basis), ascorbic acid ($C_6H_8O_6$, AA, ≥ 95%), and Nafion perfluorinated resin solution (5 wt %) were purchased from Sigma Aldrich. Cetyltrimethylammonium chloride (CTAC, 96%), cetyltrimethylammonium bromide (CTAB, > 98.0%), and citric acid monohydrate were purchased from Alfa Aesar. All reagents were used as received without further purification. Ultrapure water (18.2 MΩ resistivity, Barnstead EasyPure II 7138) was used for all experiments.

**Synthesis of CNM-Au8.** CNM-Au8 nanocrystals were synthesized following a previously published and patented protocol.[1,2] In brief, water containing 6.5 mM $NaHCO_3$ was transferred to a trough through which the buffered water flowed at a constant rate. The buffered water was exposed to an electrical plasma generated between an Au electrode, suspended above the water, and the surface of the buffered water, which functioned as a second electrode to generate plasma-conditioned water. The conditioned water was subsequently exposed to a series of paired Au wire electrodes. Each Au wire electrode pair was held by a controller that advanced the electrode pairs through the conditioned water with continuous application of alternating current as the conditioned water flowed past each electrode pair. The resulting Au nanocrystal suspension was subsequently concentrated, filtered with an antimicrobial filter, then filled into single unit high density polyethylene (HDPE) containers. All processes were executed in a clean room. Each batch of CNM-Au8 was analytically assayed as described in detail in a previous publication[2] to ensure that the released specification standards were met.

**Synthesis of CTAC-Capped Au Quasi-Spherical Nanoparticles (QSNPs).** CTAC-capped Au QSNPs with an average diameter of 11 nm were synthesized following a previously reported protocol.[3] Au seeds, around 2 nm in size, were first synthesized by injecting 1 mL of 6 mM ice-cold $NaBH_4$ into a 10 mL mixture of 0.25 mM $HAuCl_4$ and 100 mM CTAB under vigorous magnetic stir. After stirring for 2 mins, the as-obtained colloidal seeds were kept undisturbed in dark for 3 h. 50 μL of the Au seeds were added into a mixture of 1.5 mL of 100 mM AA and 2 mL of 200 mM CTAC. Then 2 mL of 0.5 mM $HAuCl_4$ were then added to initiate the nanocrystal growth. After stirring the reactant mixtures for 15 minutes, the resulting colloidal Au QSNPs were centrifuged, washed with water through three cycles of centrifugation/redispersion, and finally redispersed in water.

**Synthesis of CTAC-Capped Au Surface-Roughened Nanoparticles (SRNPs).** CTAC-capped Au SRNPs with an average diameter of 152 nm were synthesized using a seed-mediated growth method.[4-7]

The CTAC-capped colloidal Au seeds (~ 3 nm in diameter) were first synthesized by reducing $HAuCl_4$ with $NaBH_4$ in the presence of CTAC. Briefly, 0.25 mL of 10 mM $HAuCl_4$ solution was added into 10 mL of 100 mM CTAC solution under magnetic stir. Then 0.30 mL of 10 mM fresh, ice-cold $NaBH_4$ was injected into the mixture. The solution was stirred vigorously for 1 min, then left undisturbed for 2 h, and finally diluted with 100 mM CTAC by 1000-fold for future use. The growth solution for Au SRNPs consisted of 0.50 mL of 10 mM $HAuCl_4$, 0.10 mL of 100 mM AA, and 10.0 mL of 100 mM CTAC. 20 μL of the seeds were introduced into the growth solution, and the mixtures were gently mixed for 30 s, then left undisturbed for 12 h. The as-synthesized Au SRNPs were washed with water 3 times before finally being redispersed in water.

#### S1.2. Characterizations of CNM-Au8

The particle sizes and morphologies of gold nanocrystals in CNM-Au8 were characterized by transmission electron microscopy (TEM) using a Hitachi HT-7800 transmission electron microscope operated at an accelerating voltage of 100 kV. The colloidal CNM-Au8 nanocrystals were drop-dried on 300 mesh Formvar/carbon coated Cu grids (Electron Microscopy Science Inc.) for the TEM imaging. The light extinction spectra of colloidal CNM-Au8 nanocrystals were collected at room temperature using a HP-Agilent 8453 UV-Vis-NIR spectrophotometer. Electrochemical measurements were performed using a CHI 660E workstation (CH Instruments, Austin, TX) with a standard three-electrode system, with a glassy carbon electrode (GCE, 3.0 mm diameter), a saturated calomel electrode (SCE), and a platinum wire serving as the working electrode, the reference electrode, and the counter electrode, respectively. The GCE was polished with 0.3 mm alumina slurry and then washed thoroughly with water and ethanol. A colloidal ink was prepared by mixing 500 mg $L^{-1}$ CNM-Au8 with 6.5 mM of $NaHCO_3$ vehicle at volume ratio of 1:1. 10 μL of the CNM-Au8 ink were drop-casted on a pretreated GCE and dried in air at room temperature for 3 h. Then 2 μL of Nafion perfluorinated resin solution (0.2 wt %) was drop-dried on the electrode surface to hold the nanoparticles. For the electrochemical oxide stripping assays, CV measurements were conducted in $N_2$-saturated 0.5 M $H_2SO_4$ electrolyte solution at room temperature with a potential sweep rate at 5 mV $s^{-1}$. CV measurements on electro-oxidation of nicotinamide adenine dinucleotide (NADH, $C_{21}H_{27}N_7Na_2O_{14}P_2$, Bioworld, Inc.) were carried out in electrolytes containing 1.0 mM NADH in 5 mM citrate buffers at various pHs. The pH of citrate buffer was adjusted in the range 3 to 9 by varying the molar ratios of citric acid, sodium citrate, and sodium hydroxide.

#### S1.3. *In vitro* Neuroprotection Assays

**Preparation of Aβ (1-42) Oligomers**. Aβ(1-42) oligomers were prepared following a previously reported protocol.[8] Briefly, Aβ(1-42) peptide (Bachem, Weil-am-Rhein, Germany) was dissolved in the defined culture medium for primary spinal motor neurons described below, devoid of serum, at an initial concentration of 40 μM. This solution was gently agitated for 3 days at 37 °C in the dark and immediately used after being diluted in culture medium to 20 μM of Aβ(1-42) preparation containing 2 μM of Aβ oligomers (AβO), as measured by automated Western Blot.

**Preparation and Treatment of Primary Rat Spinal Cord Neurons.** Rat spinal cord MNs were cultured as described by Martinou et al.[9] and by Wang et al..[10] Briefly, embryonic day 14 (E14) spinal cords were collected from fetuses of pregnant female Wistar rats (Janvier Labs, Le Genest-Saint-Isle, France), and immediately placed in ice-cold L15 Leibovitz medium (L15, Pan Biotech, Aidenbach, Germany) with a 2% penicillin (10,000 U $mL^{-1}$), streptomycin (10 mg $mL^{-1}$) solution (PS, Pan Biotech, Aidenbach, Germany), and 1% bovine serum albumin (BSA, Pan Biotech, Aidenbach, Germany). Spinal cords were treated at 37°C for 20 min with a trypsin-EDTA solution at a final concentration of 0.05% trypsin and 0.02% EDTA (Pan Biotech, Aidenbach, Germany). The reaction was stopped by addition of Dulbecco's modified Eagle's medium (DMEM, Pan Biotech, Aidenbach, Germany) with 4.5 g/L of glucose containing 0.5 mg/mL DNAse I grade II (Pan Biotech, Aidenbach, Germany) and 10% fetal calf serum (FCS, Thermo Fisher Scientific, Cergy Pontoise, France). Cells were mechanically dissociated by three forced passages through the tip of a 10-mL pipette. Cells were then centrifuged at 180 x g for 10 min at 4°C on a layer of BSA (3.5%) in L15 medium. The supernatant was discarded, and the pellets were resuspended in a defined culture medium consisting of Neurobasal medium (Thermo Fisher Scientific, Cergy Pontoise, France) with a 2% solution of B27 supplement (Invitrogen, Carlsbad, CA, USA), 2 mM L-glutamine (Pan Biotech, Aidenbach, Germany), 2% of PS solution, and 10 ng $mL^{-1}$ of brain-derived neurotrophic factor (BDNF, Pan Biotech, Aidenbach, Germany). Viable cells were counted in a Neubauer cytometer, using the trypan blue exclusion test. The cells were seeded at a density of 20,000 per well in 96-well plates precoated with poly-L-lysine (Corning Biocoat, Le Pont de Claix, France) and cultured at 37°C in an air (95%)-$CO_2$ (5%) incubator. The medium was changed every 2 days.

On day 11 of culture, CNM-Au8 was added to a final concentration (10, 32, 100, or 316 ng $mL^{-1}$) into fresh culture medium. Spinal cord neurons were incubated with this medium for 48 hours to allow for CNM-Au8 membrane penetration. For the vehicle control, the vehicle solution (6.5 mM $NaHCO_3$ in pharmaceutical-grade water) was added to fresh media at the same volume used for the highest concentration of CNM-Au8, and this media was added to cells. 48 hours after CNM-Au8 or vehicle incubation, glutamate (5 μM, Sigma-Aldrich, Saint Quentin-Fallavier France) or prepared Aβ(1-42) oligomers (20 μM) was added to the cultures for 20 min or 24 h, respectively, in medium containing vehicle, CNM-Au8, or riluzole at the pre-treatment concentrations. After treatment with glutamate or Aβ(1-42) oligomers, cells were washed and fresh culture medium was added for an additional 48h in the presence of vehicle, CNM-Au8, or riluzole (Sigma-Aldrich). Cells were then processed for immunofluorescence or biochemical analyses as described below.

**Preparation and Treatment of Primary Rat Dopaminergic Neurons.** Rat dopaminergic neurons were cultured as described by Visanji and coworkers.[11] Briefly, the midbrains of 15-day-old rat embryos (day E15) were dissected under a microscope. The ventral portion of the mesencephalic flexure, a region of the developing brain rich in dopaminergic neurons, was used for the cell preparations. Mesencephalic tissue was processed as described above for primary spinal cord neurons. Defined culture medium for mesencephalic co-cultures consisted of Neurobasal (Invitrogen) supplemented with B27 (2%), L-glutamine (2 mM) and 2% of PS solution and 10 ng/ml of Brain-derived neurotrophic factor (BDNF) and 1 ng/ml of Glial-Derived Neurotrophic Factor (GDNF). Viable cells were counted in a Neubauer cytometer using the trypan blue exclusion test. The cells were seeded at a density of 40,000 cells/well in

96 well-plates pre-coated with poly-L-lysine and maintained in a humidified incubator at 37°C in 5% $CO_2$/95% air atmosphere. Half of the medium was changed every 2 days with fresh medium.

On day 4 of culture, CNM-Au8 was added to a final concentration (10, 32, 100, 316 ng/mL) into fresh culture medium. Mesencephalic neurons were incubated with this medium for 48 hours. For the vehicle control, the vehicle solution was added at the same volume used for the highest concentration of CNM-Au8. 6-OHDA was then added to a final concentration of 20 $\mu$M, diluted in control medium in the presence of CNM-Au8 or vehicle for 48 hours. After 6-OHDA intoxication, the mesencephalic neuronal co-culture was processed for immunofluorescence analyses as described below.

**Preparation and Treatment of Primary Rat Cortical and Hippocampal Neurons.** Rat cortical neurons were cultured as described by Callizot et al..[8] Briefly, cortices of day E15 embryos were dissected and processed as described above for primary spinal cord neurons. The defined culture medium for cortical neurons consisted of Neurobasal medium with a 2% solution of B27 supplement, 2 mM L-glutamine, 2% PS solution, and 10 ng $mL^{-1}$ BDNF. Viable cells were counted in a Neubauer cytometer, using the trypan blue exclusion test. The cells were seeded at a density of 30,000 per well in 96-well plates precoated with poly-L-lysine and were cultured at 37°C in an air (95%)-$CO_2$ (5%) incubator. The medium was changed every other day.

On day 11 of culture, CNM-Au8 was added to a final concentration (10, 32, 100, 316 ng $mL^{-1}$) into fresh culture medium. Cortical neurons were incubated with this medium for 48 hours. After 48 h, glutamate (40 µM) in the presence of vehicle or CNM-Au8 was added for 20 min. The cells were then washed, and fresh culture medium was added for an additional 48 h in the presence of vehicle or CNM-Au8. Cells were then processed for immunofluorescence or biochemical analyses as described below.

Hippocampal neurons were prepared as described for cortical neurons, with the following differences: E17 hippocampi were dissected and processed for primary culture. The cells were seeded at a density of 20,000 per well in 96-well plates precoated with poly-L-lysine.

On day 17 of culture, CNM-Au8 was added to a final concentration (10, 32, 100, 316 ng/mL) into fresh culture medium. Hippocampal neurons were incubated with this medium for 48 hours. After 48 h, glutamate (20 µM) in the presence of vehicle or CNM-Au8 was added for 20 min. The cells were then washed, and fresh culture medium was added for an additional 48h in the presence of vehicle or CNM-Au8. Cells were then processed for immunofluorescence or biochemical analyses as described below.

**Immunofluorescent Staining of Primary Cultures.** Medium was removed and cells were fixed by a cold solution of ethanol (96%) and acetic acid (5%) for 5 min at -20 °C. Cells were washed twice in phosphate buffered saline (PBS, Invitrogen, Carlsbad, CA, USA, then were permeabilized with a solution of PBS and 0.1% of saponin. PBS with1% FCS was used for blocking nonspecific antibody binding sites for 2 h at room temperature. Cultures were incubated with primary antibodies in PBS containing 1% fetal calf serum and 0.1% of saponin for 2 h at room temperature as described below.

Mouse monoclonal anti-neurofilament (NF, Sigma-Aldrich) was used at a dilution of 1/400 to label cell bodies and neurites of mature MNs. Rabbit polyclonal anti-TDP-43 (Cell signaling, Ozyme, Saint-Cyr-l'Ecole, France) was used at a dilution of 1/100 to label both nuclear and cytoplasmic TDP43. Nuclear

TDP-43 was identified by overlap of signal with Hoechst stain (Sigma-Aldrich), which was used at 1/1000 in the same solution with anti-TDP-43 antibodies. Mouse monoclonal anti-Tyrosine Hydroxylase (TH, Sigma-Aldrich) was used at a dilution of 1/10,000 to label dopaminergic cell bodies and neurites. Polyclonal rat anti-α-synuclein was used at the dilution of 1/200 (Cell Signaling). Mouse monoclonal anti microtubule-associated-protein 2 (MAP-2, Sigma-Aldrich) was used at dilution of 1/400 to label hippocampal and cortical cell bodies and neurites. Rabbit polyclonal anti-PSD95 (post synaptic density protein 95, Abcam, Cambridge, United Kingdom) was used at dilution of 1/400.

Secondary antibodies used were Alexa Fluor 488 goat anti-mouse IgG (Thermo Fisher Scientific, Illkirch-Graffenstaden, France) at the dilution of 1/400-1/800, and Alexa Fluor 568 goat anti-rabbit IgG (Thermo Fisher Scientific, Illkirch-Graffenstaden, France) at the dilution of 1/400. Appropriate secondary antibodies were diluted in PBS containing 1% FCS, 0.1 % saponin, and cells were incubated for 1 h at room temperature.

**Imaging Neuronal Cultures.** Images were acquired with a confocal microscope LSM 900 with Zen software. For each condition, 20-30 pictures per well (representing ≥90 % of the well area) were automatically taken using ImageXpress (Molecular Devices, LLC, San Jose, CA) with 10x-20x magnification. All images were taken with the same conditions. Analysis was performed automatically by using Custom Module Editor (Molecular Devices).

***In vitro* Quantitation of $NAD^+$ and NADH.** Primary rat mesencephalic neural-glia co-cultures were seeded at a density of 40,000 cells/well in 96-well plates pre-coated with poly-L-lysine. After four days of culture, CNM-Au8 (10 ng $mL^{-1}$, 100 ng $mL^{-1}$, 1 $\mu$g $mL^{-1}$, 10 $\mu$g $mL^{-1}$), BDNF (50 ng/mL), or vehicle was added for 36 h. Quantitation of $NAD^+$ and NADH was performed by bioluminescent assay (Promega kit #G9071).

***In vitro* Quantitation of Adenosine Triphosphate (ATP).** M03.13 cells (TebuBio) were seeded at a density of 20,000 cells/well in a 96-well plate. After one day of culture, cells were treated with CNM-Au8 (0.1 $\mu$g/mL, 0.32 $\mu$g/mL, 1 $\mu$g/mL) or vehicle for 72 h. Half of the wells (N = 3 per condition) were treated with mitochondrial blockers (100 $\mu$M antimycin A, 0.5 $\mu$M rotenone, 3 $\mu$M oligomycin) for two hours. Cells were then lysed in luciferase buffer. ATP was determined by bioluminescence assay (1 s integration time, CLARIOstar). Mitochondrial ATP was calculated by subtracting the average ATP level from mitochondria-blocked cells from average total ATP levels of untreated cells.

**Statistical Analyses of *in vitro* Experiments.** All values are expressed as mean ± SEM (standard error of the mean) from 6 wells per condition per culture. Data is expressed as a percentage of control conditions, with the no injury, vehicle-treated control set at 100 %. Statistical analyses using GraphPad Prism software version 9.5.1 (GraphPad Software, LLC, La Jolla, CA, USA) using one-way ANOVA followed by Dunnett's test or PLSD Fisher's test, as appropriate, to correct for multiple comparisons.

### S2. Additional Figures

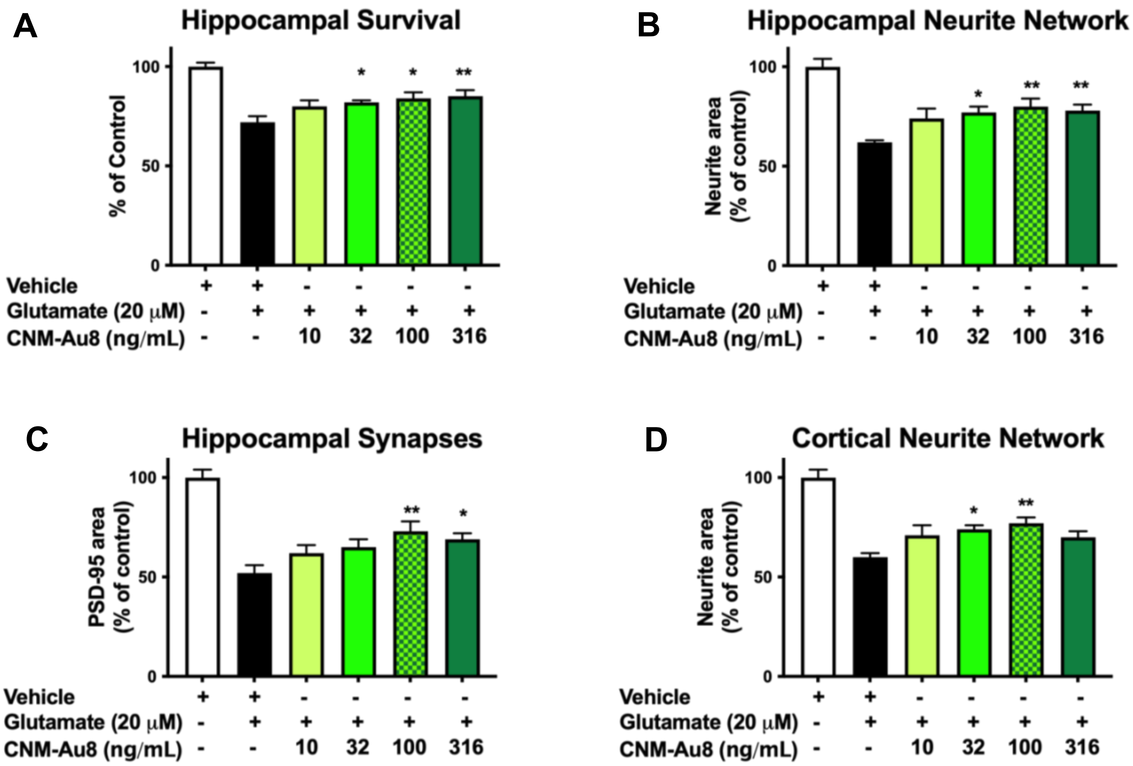


***Figure S1.*** CNM-Au8 treatment increases hippocampal neuronal survival and protects hippocampal and cortical neurite networks from glutamate challenge in vitro. Primary rat E17 hippocampal neurons (A-C) or cortical neurons (D) were cultured for 17 days, and then treated with vehicle or CNM-Au8. Glutamate (20 mM) was added to the cultures for 20 min to induce neuronal death. After 48h, cultures were fixed and stained with anti-MAP2 to quantitate neuron survival (A) and neurite length (B, D), and co-stained with anti-PSD95 to quantitate intact synapses (C). Six replicates were performed per condition. Group means +/- SEM. * $p < 0.05$; ** $p < 0.01$; treatment vs. vehicle, one way ANOVA corrected for multiple comparisons.

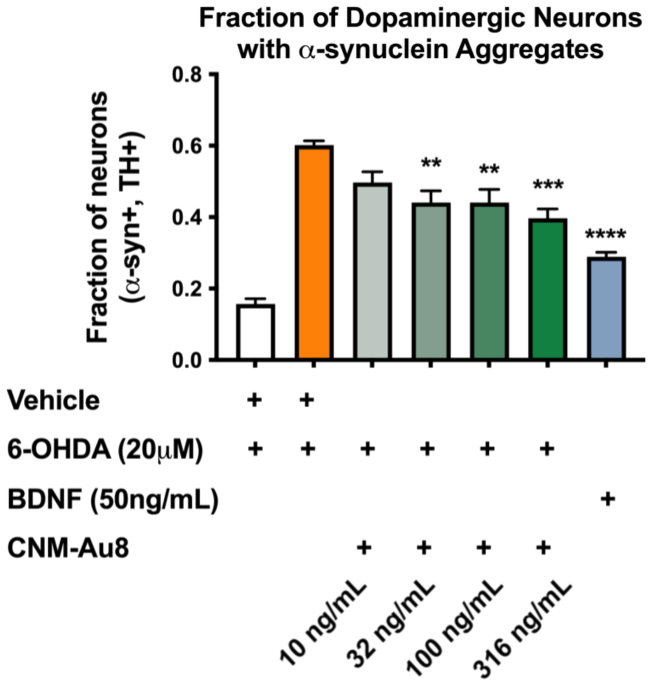


***Figure S2.*** CNM-Au8 treatment reduces the number of dopaminergic neurons that accumulate cytoplasmic α-synuclein aggregates in response to the neurotoxin 6-hydroxydopamine (6-OHDA). Primary rat E15 dopaminergic neurons were cultured for four days then treated with vehicle, CNM-Au8, or 50 ng/mL BDNF (positive control). 6-OHDA (20 mM) was added to induce accumulation of a-synuclein aggregates. After 48h, cells were fixed and co-stained with anti-tyrosine hydroxylase antibodies (TH) to mark dopaminergic cells and anti-alpha-synuclein antibodies. The fraction of α-syn positive, TH-positive neurons out of the total TH-positive neurons was quantitated by automated system. Six replicates were performed per condition. Group means +/- SEM. * $p < 0.05$; ** $p < 0.01$; *** $p < 0.001$; **** $p < 0.0001$; treatment vs. vehicle, one way ANOVA corrected for multiple comparisons.

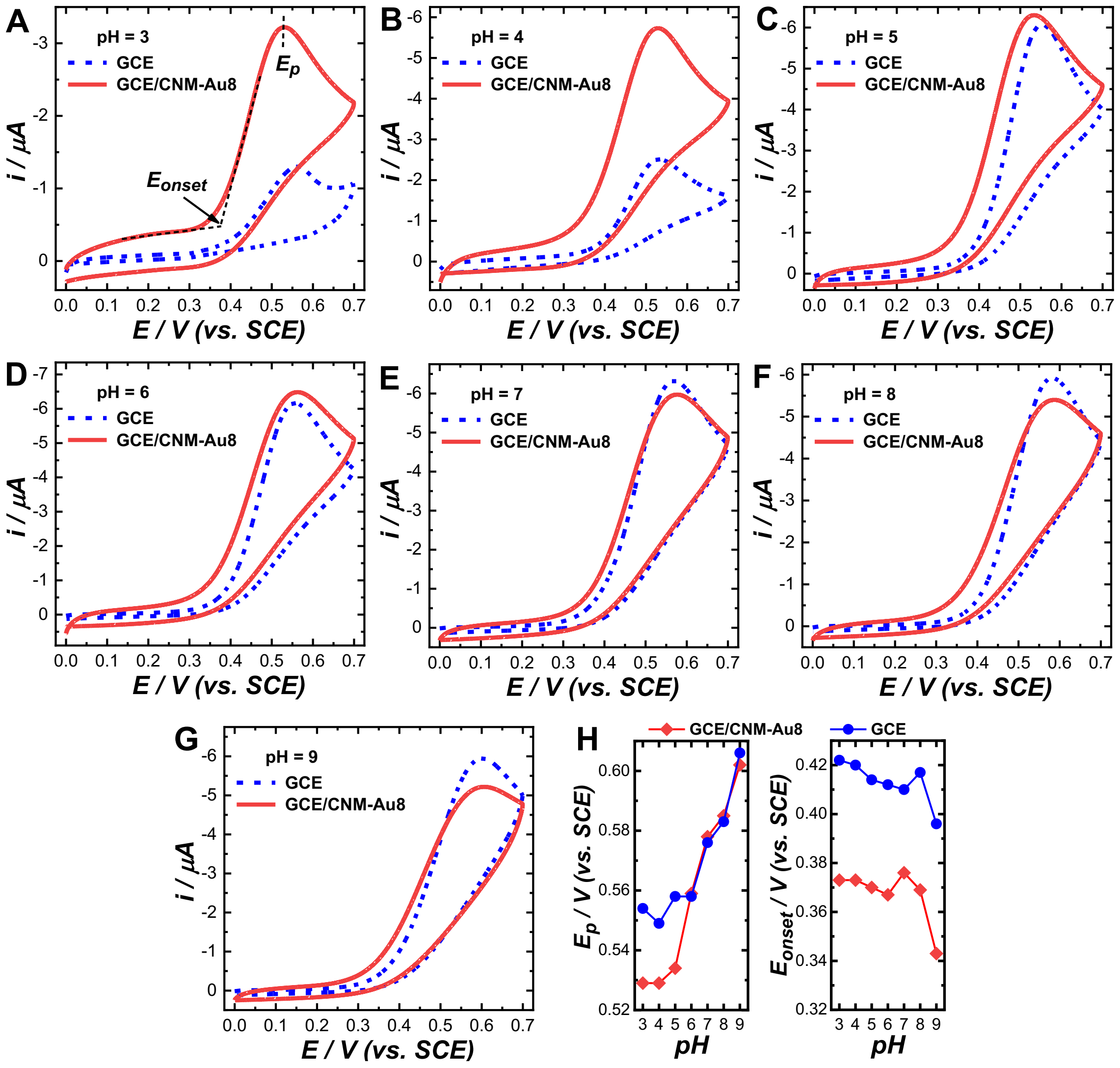


***Figure S3.*** Cyclic voltammetry curves collected from electrolytes containing 1 mM NADH at pHs of (A) 3, (B) 4, (C) 5, (D) 6, (E) 7, (F) 8, and (G) 9. The curves collected when a bare glassy carbon electrode (GCE) served as the working electrode were shown as dash blue curves and those collected on CNM-Au8-loaded GCEs (GCE/CNM-Au8) were shown as red solid curves. (H) Peak potentials, $E_p$, and onset potentials, $E_{onset}$, at various pHs. A saturated calomel electrode (SCE) and a platinum wire were used as the reference electrode and the counter electrode, respectively, in the electrochemical measurements. In the electrolytes, 1 mM NADH was dissolved in 5 mM citrate buffers at various pHs. The pH of citrate buffer was adjusted in the range 3 to 9 by varying molar ratios of citric acid monohydride, sodium citrate dihydrate and sodium hydroxide.

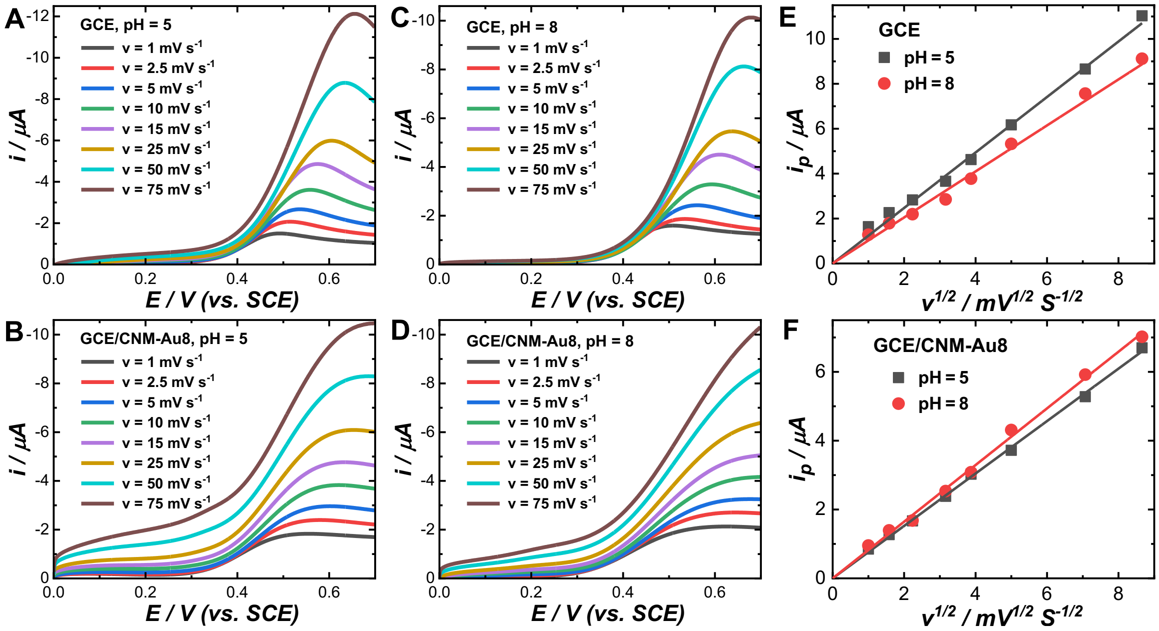


***Figure S4.*** Linear sweep voltammetry curves of electro-oxidation of NADH at a pH of 5 on (A) GCE and (B) GCE/CNM-Au8 and at a pH of 8 on (C) GCE and (D) GCE/CNM-Au8 at various potential sweep rates. The concentration of NADH in the electrolytes was 1 mM. Relationships between peak current, $i_p$, and square root of potential sweep rate, $v^{1/2}$, at pHs of 5 and 8 on (E) GCE and (F) GCE/CNM-Au8.

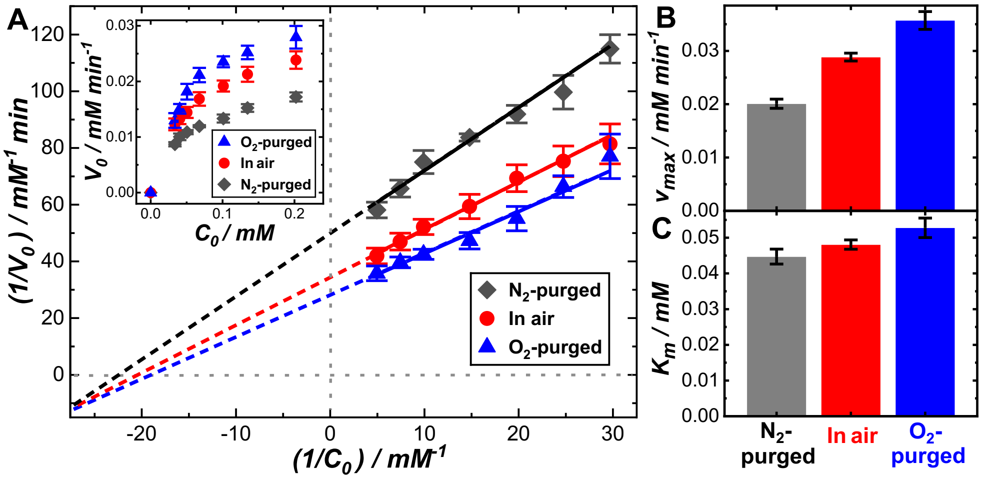


***Figure S5.*** (A) Lineweaver-Burk plots showing the relationships between the reciprocal of initial velocity and the reciprocal of initial NADH concentration for the reactions occurring under ambient air and in $N_2$-purged or $O_2$-purged solutions. The initial concentration of NADH was 0.10 mM and the total volume of the catalyst-reactants mixtures was kept at 2.0 mL. The mass concentration of CNM-Au8 was 37.7 mg $L^{-1}$. The reactions occurred at room temperature. The relationships between the initial velocity and initial NADH concentration are shown in the inset. Comparison of the values of (B) $v_{max}$ and (C) $K_m$ in $N_2$-purged, $O_2$-purged, and ambient environments. The pH value was maintained at 5 for all reactions. The errors bars in panel A represent the standard deviations of 3 experimental runs. The error bars in panels B and C represent the standard deviations associated with least squares curve fitting.

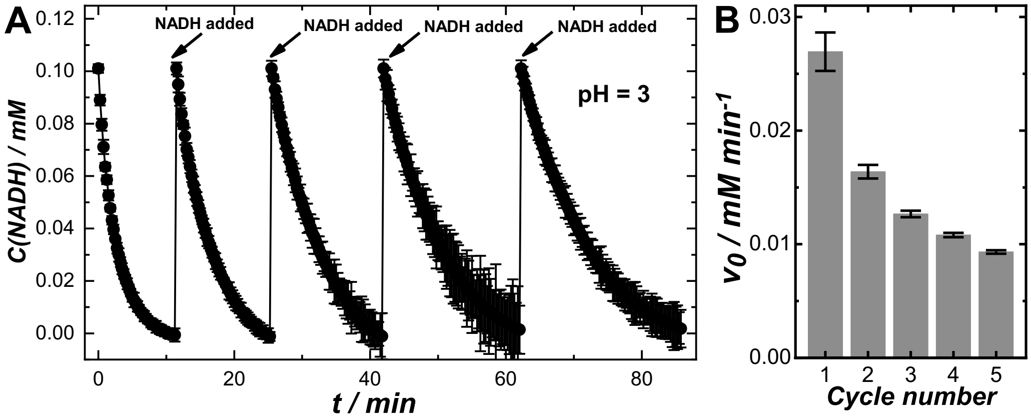


***Figure S6.*** (A) Temporal evolution of NADH concentration at a pH of 3 over multiple reaction cycles. In each reaction cycle, the initial concentration of NADH was 0.10 mM and the total volume of the catalyst-reactants mixtures was kept at 2.0 mL. The mass concentration of CNM-Au8 was 37.7 mg $L^{-1}$. After NADH oxidization reactions approached completion in each cycle, 2 $\mu$L of 100 mM NADH was added to bring the NADH concentration back to 0.10 mM to initiate the next reaction cycle. The time spots at which NADH was added were labeled with arrows. The reactions occurred at room temperature. (B) Evolution of initial velocity over multiple reaction cycles. The errors bars in panel A represent the standard deviations of 3 experimental runs. The error bars in panel B represent the standard deviations associated with least squares curve fitting.

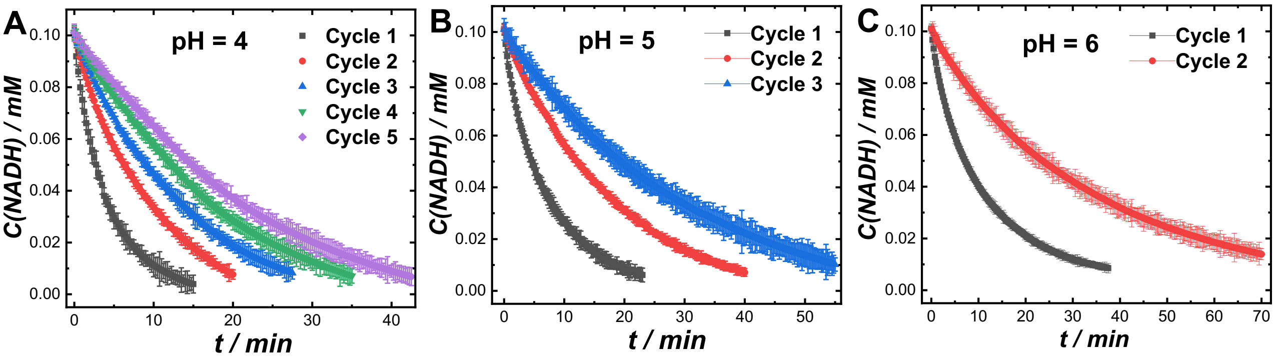


***Figure S7.*** (A) Temporal evolutions of NADH concentration at pHs of (A) 4, (B) 5, and (C) 6 over multiple reaction cycles. In each reaction cycle, the initial concentration of NADH was 0.10 mM and the total volume of the catalyst-reactants mixtures was kept at 2.0 mL. The mass concentration of CNM-Au8 was 37.7 mg $L^{-1}$. After NADH oxidization reactions approached completion in each cycle, 2 $\mu$L of 100 mM NADH was added to bring the NADH concentration back to 0.10 mM to initiate the next reaction cycle. The reactions occurred at room temperature. The errors bars represent the standard deviations of 3 experimental runs.

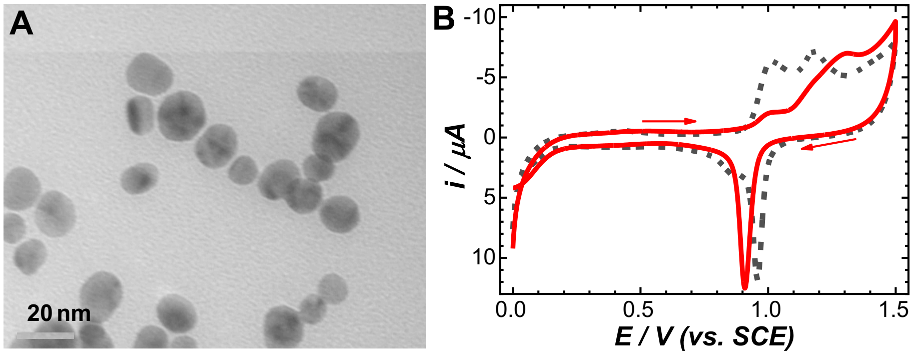


***Figure S8.*** (A) Transmission electron microscopy (TEM) image of CNM-Au8 recycled after 5 reaction cycles at a pH of 3. (B) Cyclic voltammetry curves collected from as-synthesized CNM-Au8 (dash grey curve) and CNM-Au8 recycled after 5 reaction cycles (solid red curve) in an $N_2$-purged aqueous electrolyte containing 0.5 M $H_2SO_4$ at room temperature. The potential sweep rate was 5.0 mV $s^{-1}$ and the arrows showed the direction of potential sweep.

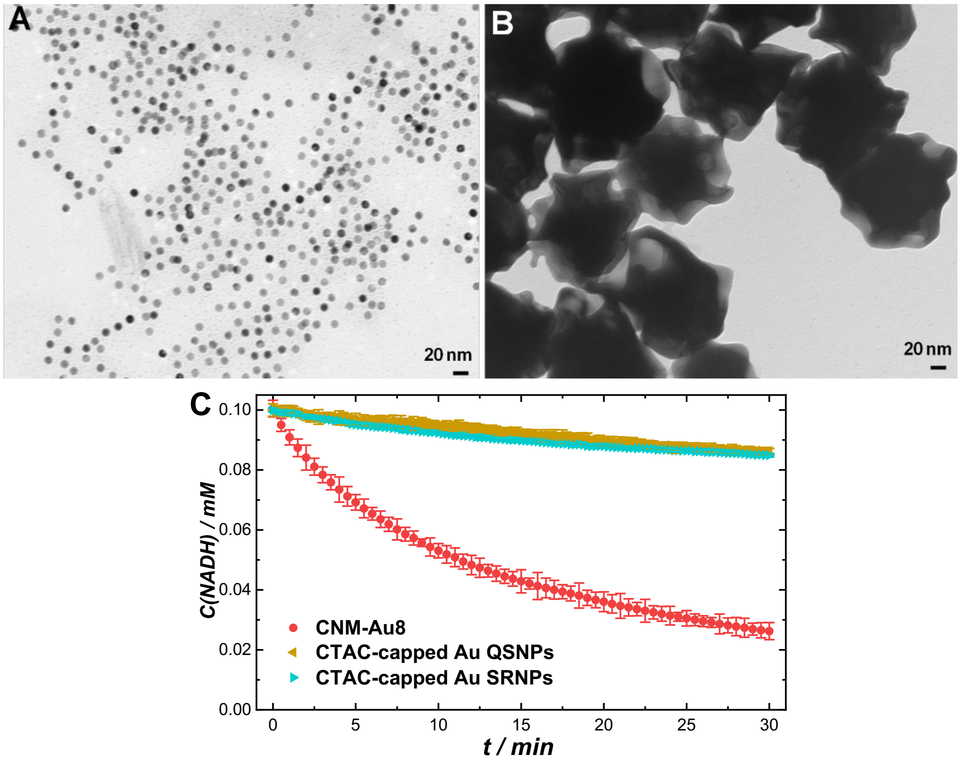


***Figure S9.*** TEM images of (A) cetyltrimethylammonium chloride (CATC)-capped Au quasi-spherical nanoparticles (QSNPs) and (B) CTAC-capped Au surface-roughened nanoparticles (QSNPs). (C) Temporal evolutions of NADH concentration during NADH oxidation reactions catalyzed by CNM-Au8, CATC-capped Au QSNPs, and CTAC-capped Au SRNPs at a pH of 7 at room temperature. In each reaction cycle, the initial concentration of NADH was 0.10 mM and the total volume of the catalyst-reactants mixtures was kept at 2.0 mL.

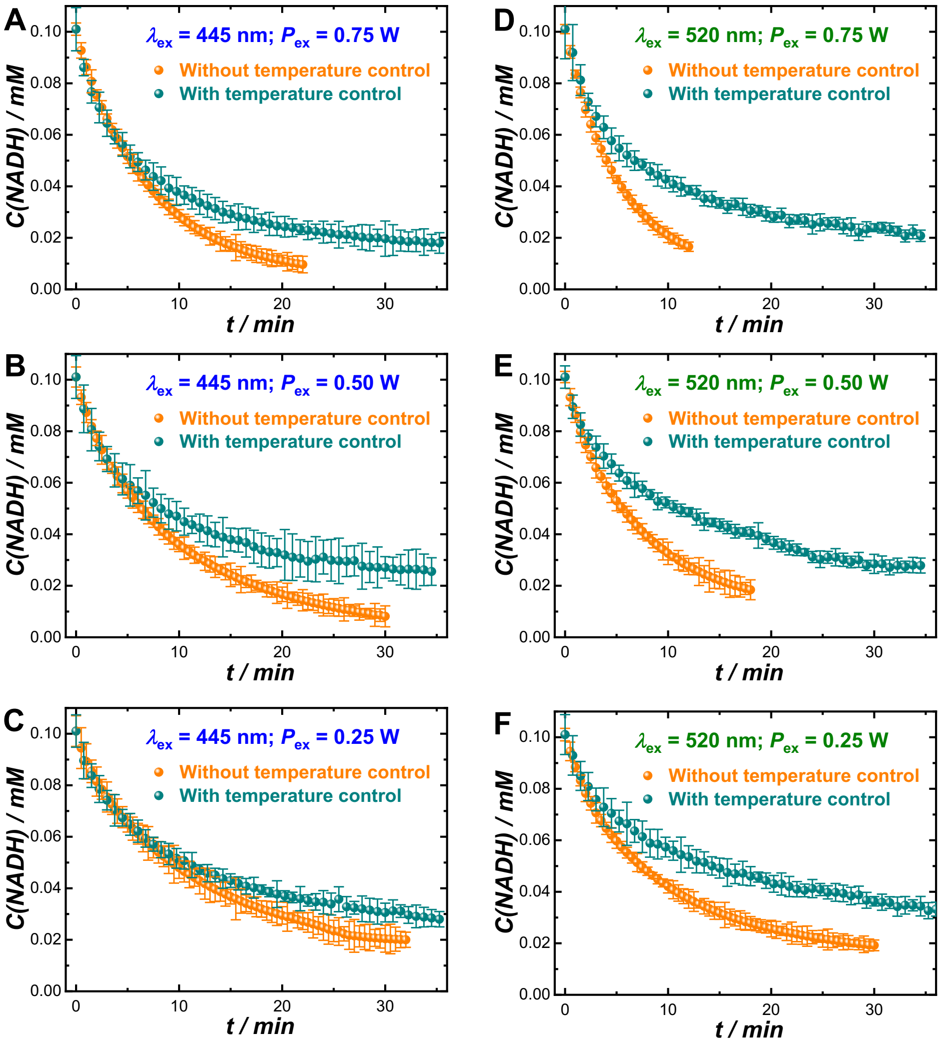


***Figure S10.*** Temporal evolutions of NADH concentration during CNM-Au8-catalyzed NADH oxidation reactions under photo-illumination at $\lambda_{ex}$ of 445 nm and $P_{ex}$s of (A) 0.75, (B) 0.50, and (C) 0.25 W and at $\lambda_{ex}$ of 520 nm and $P_{ex}$s of (D) 0.75, (E) 0.50, and (F) 0.25 W with and without temperature control. In all reactions, the mass concentration of Au catalysts was kept at 37.7 mg $L^{-1}$ and the initial concentration of NADH was 0.10 mM. The reactions occurred at a pH of 8. The errors bars represent the standard deviations of 3 experimental runs.

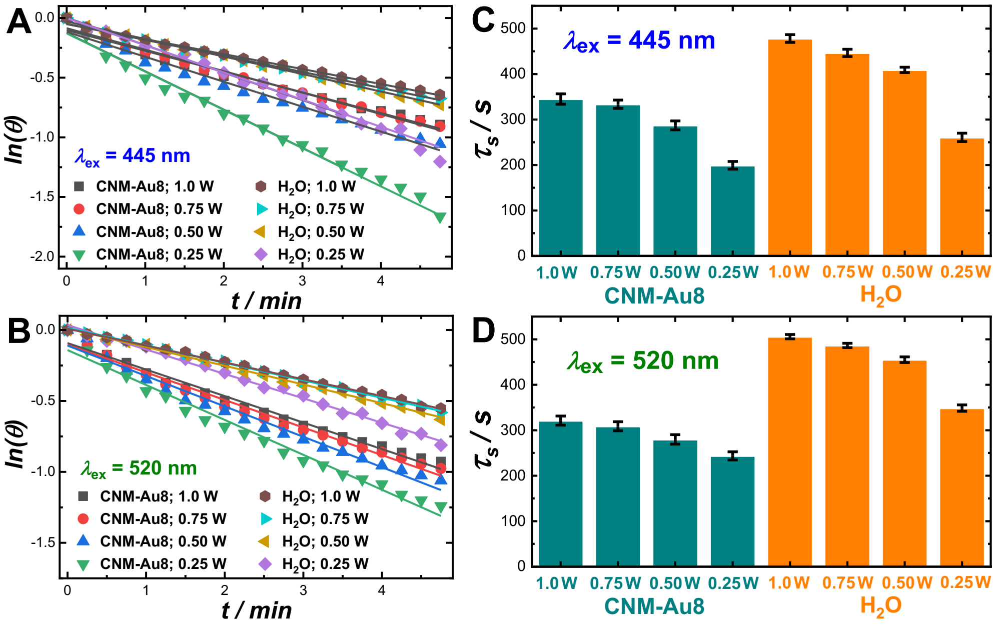


***Figure S11.*** Temporal evolution of ln($\theta$) during the first 5 minutes in the cooling processes for colloidal CNM-Au8 suspensions (Au mass concentration of 37.7 mg L$^{-1}$) and pure water following laser illuminations at $\lambda_{ex}$s of (A) 445 and (B) 520 nm and various $P_{ex}$s. $\tau_s$ values obtained from least squares curve fitting for CNM-Au8 and water at various $P_{ex}$s at $\lambda_{ex}$s of (C) 445 and (D) 520 nm.

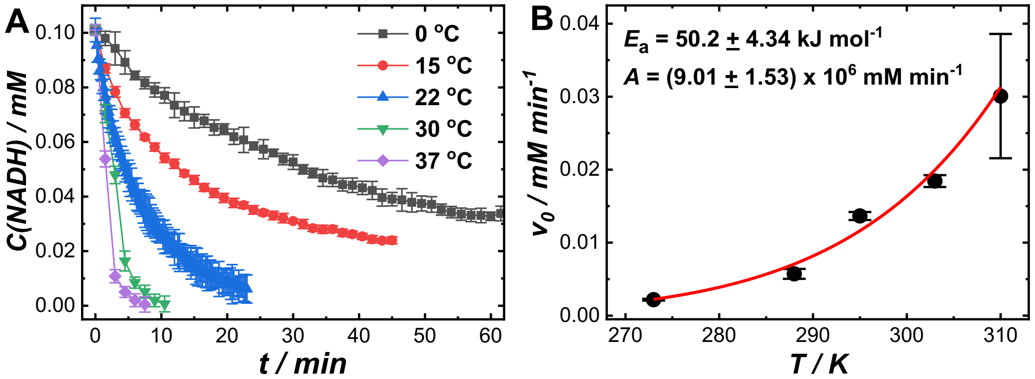


***Figure S12.*** (A) Temporal evolutions of NADH concentration during CNM-Au8-catalyzed NADH oxidation reactions in dark at a pH of 5 at various reaction temperatures. (B) Relationship between $v_0$ and reaction temperature, $T$, at a pH of 5. The experimental results were fitted with the Arrhenius equation. In all reactions, the mass concentration of Au catalysts was kept at 37.7 mg L$^{-1}$ and the initial concentration of NADH was 0.10 mM. The errors bars in panel A represent the standard deviations of 3 experimental runs. The error bars in panel B represent the standard deviations associated with least squares curve fitting.
